# Quantifying the individual impact of artificial barriers in freshwaters: A standardized and absolute genetic index of fragmentation

**DOI:** 10.1101/809111

**Authors:** Jérôme G. Prunier, Camille Poesy, Vincent Dubut, Charlotte Veyssière, Géraldine Loot, Nicolas Poulet, Simon Blanchet

## Abstract

Fragmentation by artificial barriers is an important threat to freshwater biodiversity. Mitigating the negative aftermaths of fragmentation is of crucial importance, and it is now essential for environmental managers to benefit from a precise estimate of the individual impact of weirs and dams on river connectivity. Although the indirect monitoring of fragmentation using molecular data constitutes a promising approach, it is plagued with several constraints preventing a standardized and individual quantification of barrier effects. Indeed, observed levels of genetic differentiation depend on both the age of the obstacle and the effective size of the populations it separates, making difficult comparisons of the actual barrier effect of different obstacles. Here, we developed a standardized genetic index of fragmentation (*F*_*INDEX*_), allowing an absolute and independent assessment of the individual effects of obstacles on connectivity. The *F*_*INDEX*_ is the standardized ratio (expressed as a percentage) between the observed genetic differentiation between pairs of populations located on either side of an obstacle and the genetic differentiation expected if this obstacle completely prevented gene flow. The expected genetic differentiation is calculated from simulations taking into account two nuisance parameters: the number of generations since barrier creation (the age of the obstacle) and the expected heterozygosity of the targeted populations, a proxy for effective population sizes. Using both simulated and published empirical datasets, we explored and discussed the validity and the limits of the *F*_*INDEX*_. We demonstrated that it allows quantifying genetic effects of fragmentation only from a few generations after barrier creation and provides valid comparisons among populations (or species) of different effective populations sizes and obstacles of different ages. The computation of the *F*_*INDEX*_ requires a minimum amount of fieldwork and genotypic data, and solves some of the difficulties inherent to the study of artificial fragmentation in rivers and potentially in other ecosystems. This makes the *F*_*INDEX*_ a promising and objective tool for managers aiming at at planning restoration programs and at evaluating the efficiency of these programs.

## Introduction

Heavily impacted by human activities, rivers are at the heart of biodiversity conservation issues (Dudgeon et al., 2006; Reid et al., 2018). Among the various threats to these ecosystems, river fragmentation by artificial barriers is considered as the most widespread and worrying (Couto & Olden, 2018; Nilsson, 2005; Turgeon, Turpin, & Gregory-Eaves, 2019). Weirs and dams, but also pipes and culverts, have long been, and are still, constructed for flow regulation and/or hydropower supply but they often imply a loss of habitat and a reduction in riverscape functional connectivity (that is, species-specific) in freshwater organisms (Birnie-Gauvin, Aarestrup, Riis, Jepsen, & Koed, 2017; Jansson, Nilsson, & Malmqvist, 2007). For fish, artificial fragmentation is known to impact key biological processes such as migration, dispersal and recruitment, and thus viability and productivity of populations and communities (Blanchet, Rey, Etienne, Lek, & Loot, 2010; Poulet, 2007; Turgeon et al., 2019). Given the central role of hydropower as a source of energy, mitigating these negative aftermaths is now of high importance (Couto & Olden, 2018; Gibson, Wilman, & Laurance, 2017). Different restoration and mitigation measures may be considered to enhance longitudinal river connectivity, including the removal of obstacles, periodic turbine shutdowns and fishpasses setting (Bednarek, 2001; Poff & Schmidt, 2016; Silva et al., 2018). However, these measures may all result in unintended outcomes (e.g., McLaughlin et al., 2013), or unsatisfactory trade-offs between conservation of biodiversity, preservation of historical and cultural legacy and the maintenance of services provided by obstacles (Gibson et al., 2017; Hand et al., 2018; Roy et al., 2018; Song et al., 2019). In terms of conservation planning, it is therefore essential that environmental managers benefit from precise estimates of the actual impacts of different obstacles on river connectivity, or from precise estimates of the gain in connectivity resulting from restoration actions, in order to guide the prioritization of conservation efforts and to evaluate their efficiency (Cooke & Hinch, 2013; Januchowski-Hartley, Diebel, Doran, & McIntyre, 2014; Raeymaekers, Raeymaekers, Koizumi, Geldof, & Volckaert, 2009).

The direct monitoring methods conventionally used in rivers to quantify the functional permeability of an obstacle or the efficiency of a restoration action are video-counting, telemetry and capture-recapture protocols. Although efficient (e.g., Cooke & Hinch, 2013; Hawkins et al., 2018; Junge et al., 2014; Pracheil et al., 2015), these methods are associated with technical constraints. In particular, ecological studies based on video counting or telemetry are often conducted on a limited number of obstacles, whereas robust capture-recapture protocols imply repeated and exhaustive capture sessions, ideally over several years, which involves the mobilization of substantial human and financial resources (Cayuela et al., 2018).

Indirect monitoring based on molecular data constitutes a promising alternative approach, allowing multi-specific studies of dam-induced fragmentation (Selkoe, Scribner, & Galindo, 2015). Among the many analytical procedures developed in recent years to quantify the mobility of organisms on the basis of genetic or genomic data, assignment methods and parentage analyses (Jombart, Devillard, & Balloux, 2010; Pritchard, Stephens, & Donnelly, 2000; Städele & Vigilant, 2016; Wilson & Rannala, 2003) allow the detection of =real-time’ non-effective movements (that is, not necessarily followed by a reproduction event; e.g., Junge et al., 2014; Raeymaekers et al., 2009; Saint-Pé et al., 2018) but they require an extensive sampling of individuals and moderate to high genetic differentiation between populations (Broquet & Petit, 2009; Cayuela et al., 2018).

An alternative method to quantify the permeability of an obstacle from molecular data is simply to measure the level of neutral genetic differentiation between populations located in the immediate upstream and downstream vicinity of an obstacle (i.e., located a few hundreds of meters to one kilometer apart, an *adjacent sampling strategy*), an approach that does not necessarily require large sample sizes (i.e., *n* ∼ 20-30) or heavy computation: any drop in local functional connectivity due to the creation of a barrier to gene flow is expected to translate into an increase in neutral genetic differentiation (Raeymaekers et al., 2009). However, measures of neutral genetic differentiation may only be considered as correct estimates of actual barrier effects when comparing obstacles of the *same* age (in terms of number of generations since barrier creation) and/or separating populations of *similar* effective size. This is because genetic differentiation primarily stems from genetic drift, that is, from the random fluctuation of allelic frequencies naturally occurring in all populations (Allendorf, 1986). When populations are separated by an obstacle to gene flow, these fluctuations tend to occur independently in each population, leading to a differential distribution of allelic frequencies on either side of the barrier. However, this process is progressive, taking place over several generations (Landguth et al., 2010), and is all the more slow as effective population sizes are large (Broquet & Petit, 2009; Cayuela et al., 2018; Prunier, Dubut, Chikhi, & Blanchet, 2017). As a consequence, it is impossible to attribute the differences in levels of genetic differentiation observed between obstacles varying in age and/or in the effective size of populations they separate to differences in their actual barrier effects; older obstacles or obstacles separating smaller populations should show higher genetic differentiation than more recent obstacles or obstacles separating larger populations, despite similar actual barrier effects. Given this drawback, there is an urgent need for the development of a *standardized* and *absolute* genetic index of fragmentation that takes into account the contribution of both the age of the obstacle (expressed in the number of generations since barrier creation) and the effective size of populations (or a proxy of it since this parameter is notoriously difficult to quantify; Wang, 2005) to observed measures of genetic differentiation. Such an index might allow a quick and robust quantification of individual and actual barrier effects whatever their characteristics, paving the way for informed management prioritization and proper evaluation of restoration measures, along with inter-basins and interspecific comparative studies.

Here, we bridge that gap by developing a user-friendly and standardized genetic index of fragmentation (see Appendix S1 for a walkthrough), allowing an absolute and independent assessment of the individual effects of obstacles on gene flow. The proposed index (*F*_*INDEX*_) is expressed as a percentage and directly quantifies the relative loss of gene flow resulting from the presence of an obstacle. It is based on the comparison of measures of genetic differentiation observed between populations located in the immediate upstream and downstream vicinity of a putative obstacle with the theoretical measures of genetic differentiation that would be expected if the obstacle was a total barrier to gene flow. These theoretical measures of genetic differentiation are inferred from numerous genetic simulations, here used to reflect the expected changes in allelic frequencies resulting from the interplay between the age of the obstacle and the effective population sizes: the closer the observed measure of genetic differentiation from the one that would be expected in the worst-case scenario (total barrier to gene flow), the higher the index of fragmentation. We first present the logic and principles underlying our index. We then use both simulated and published empirical genetic datasets to explore and discuss the validity and the limits of the proposed index. We finally propose several perspectives to use the index and, because setting bio-indicators takes time, we present potential improvements that should be considered to make this index even more useful to managers.

## Material and methods

### Principle of the proposed genetic index of fragmentation *F*_*INDEX*_

The proposed genetic index of fragmentation *F*_*INDEX*_ is designed as a standardized estimate of the reduction in gene flow between two adjacent populations separated by an obstacle. It simply consists in rescaling the observed measure of genetic differentiation *GD*_*obs*_within its theoretical range of variation, taking into account the expected temporal evolution of allelic frequencies resulting from the interplay between the age of the obstacle and the averaged effective size of populations. This theoretical range of variation spans from *GD*_*min*_ to *GD*_*max*_ *GD*_*min*_ stands for the theoretical measure of genetic differentiation that would be expected if the obstacle was totally permeable to gene flow (crossing rate *m* ≈ 0.5). *GD*_*min*_ should theoretically equal 0 but the background noise resulting from the concomitant influences of genetic drift, mutations and incomplete genetic sampling may actually lead to non-null –though very low– measures of genetic differentiation. On the other hand, *GD*_*max*_ stands for the theoretical measure of genetic differentiation that would be expected under the worst-case scenario, that is, under the hypothesis that the considered obstacle is a total barrier to gene flow (*m* = 0). *GD*_*max*_ is expected to increase with time since barrier creation and to decrease with the increase in effective population sizes (Gauffre, Estoup, Bretagnolle, & Cosson, 2008; Landguth et al., 2010). The genetic index of fragmentation *F*_*INDEX*_ is then computed as follows (see Appendix S2 for details):

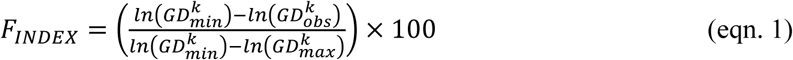

The *F*_*INDEX*_ ranges from 0 % (the observed measure of genetic differentiation is minimum-but not null- and equals the expected value *GD*_*min*_ under the assumption that the considered obstacle has no impact on gene flow) to 100 % (the observed measure of genetic differentiation is maximum and equals the expected value *GD*_*max*_ under the assumption that the considered obstacle acts as a total barrier to gene flow). The *F*_*INDEX*_ thus directly quantifies the loss of gene flow resulting from the presence of an obstacle.

*GD*_*obs*_ is directly calculated from observed genotypic data collected in populations located at the immediate upstream and downstream vicinity of the obstacle (a few hundred of meters to one kilometer apart depending on the target species; see below), whereas *GD*_*min*_ and *GD*_*max*_ are predicted from theoretical datasets simulated according to three main parameters (see the next section for details): the mutation rate *µ* of considered genetic markers, and (for *GD*_*max*_ only) the age T of the total barrier to gene flow (expressed in number of generations since barrier creation; Landguth et al., 2010; Lowe & Allendorf, 2010) and the averaged expected heterozygosity *He* of the two considered populations (a proxy for mean effective population size; see below and Appendix S5; Hague & Routman, 2016; Prunier et al., 2017).

### Expected measures of genetic differentiation

We used QuantiNemo2 (Neuenschwander, Michaud, & Goudet, 2019), an individual-based simulator for population genetics, to simulate theoretical datasets that will in turn be used to predict *GD*_*min*_ *GD*_*max*_ and values. We designed a very simple meta-population composed of two adjacent demes of carrying capacity *K*, with *K* ranging from 30 to 2000 individuals and kept constant over time. We used forward simulations of gene flow between these two demes over 1000 non-overlapping generations. Genetic polymorphism was based on 15 microsatellite loci and 20 alleles per locus, which corresponds to the number of markers typically used in empirical study focusing on functional connectivity (e.g., Blanchet et al., 2010; Coleman et al., 2018; Storfer, Murphy, Spear, Holderegger, & Waits, 2010). The mutation rate *µ*, following a stepwise mutation model, was set to 5×10^−5^ or 5×10^−4^, so as to explore the natural variability observed in microsatellite markers (mutation rate ranging from 10^−6^ to 10^−2^; Li, Korol, Fahima, Beiles, & Nevo, 2002; Schlötterer, 2000; Yue, David, & Orban, 2007). Genotypes were randomly assigned to individuals at the beginning of simulations. The inter-deme migration rate was set to 0.5 for the first 400 generations, a value providing an optimal mixing of populations and mimicking a natural situation without barrier, and then dropped to zero for the last 600 generations, mimicking the creation of a total barrier to gene flow, splitting a “single” population into two adjacent subpopulations. With populations being isolated for 600 generations, we made sure our simulations covered a realistic time frame: most artificial barriers in freshwater ecosystems were indeed built between the Middle Ages (weirs from the 12th–15th centuries) and today (largest dams built from the ∼1940’s to now), which corresponds to a number of generations ranging from 0 to ∼ 400 in most aquatic organisms such as fish species (assuming a generation time of 2 years for fish species). For each carrying capacity *K* (93 levels) and each mutation rate *µ* (2 levels), we ran ten simulation replicates, and 30 genotypes were sampled every ten generations from generation 300 to generation 1000 (71 levels) to monitor the setting up of genetic differentiation over time. This procedure resulted in a total of 93×2×71×10=132060 simulated genetic datasets in the *Fstat* format (Goudet, 1995), further converted into the *genepop* format (Rousset, 2008) using R (R Development Core Team, 2014).

For each simulation, we computed the two following pairwise metrics of genetic differentiation: the Hedrick’s G’’st (Hedrick, 2005; Meirmans & Hedrick, 2011) and the Meirmans’ φ’st (Meirmans, 2006), both computed using the R-package *mmod* (Winter, 2012). Nine other metrics were initially considered, but preliminary analyses revealed that some were dependent on sample size (e.g., the proportion of shared alleles or the Cavalli-Sforza and Edwards’ Chord distance; Bowcock et al., 1994; Cavalli-Sforza & Edwards, 1967; see Appendix S3 for details), while others were sensitive to mutation rate and/or did not show enough variability (e.g., the Weir and Cockerham’s θst or the Jost’s D; Jost, 2008; Weir & Cockerham, 1984; see Appendix S4 for details): they were thus discarded to avoid jeopardizing the validity of the proposed index. We found that the two retained metrics G’’st and φ’st were robust to variations in mutation rate and increased quickly after barrier creation, especially in the case of small effective population sizes (Appendix S4), in accordance with theoretical expectations (Lowe & Allendorf, 2010; Meirmans & Hedrick, 2011). All negative G’’st and φ’st values were set to 0. For each simulated dataset, we also computed the averaged expected heterozygosity *He* over the 15 loci in each population. *He* was then averaged over the two populations and further considered as a proxy for effective population sizes (see Appendix S5a). *He* indeed increased monotonically with carrying capacity in our simulations, in accordance with both theoretical and empirical works indicating that genetic diversity should increase with the increase in effective population sizes (Hague & Routman, 2016; Kimura, 1983). We here focused on mean heterozygosity because, unlike metrics such as allelic richness, heterozygosity values are bound between 0 and 1, which facilitates comparison between case studies. Moreover, this metric is much more straightforward to calculate for managers than the actual effective population size, since the latter is notoriously difficult to estimate in complex landscapes (Paz-Vinas et al., 2013; Wang, 2005). Note also that the use of two different realistic mutation rates yielded two levels of *He* across simulations (a low level at the low mutation rate and a high level at the high mutation rate; Appendix S5b), thus mimicking uncertainty in our proxy for effective population sizes. In addition to the two metrics of genetic differentiation G’’st and φ’st and to the expected heterozygosity *He*, we also kept record of the simulation replicate number, the mutation rate *µ*, the generation *t* at which genotypes were collected, the age T of the barrier (computed as and expressed in number of generations since barrier creation) and the carrying capacity *K* of simulated populations.

The 111600 simulations associated with T > 0 (i.e., after the creation of the barrier) were used as a training set in the regression implementation of a Random Forest machine-learning algorithm (Breiman, 2001). This approach was chosen as it is currently one of the most efficient statistical techniques for making predictions from non-linear data, with only a few parameters to tune (Genuer, Poggi, Tuleau-Malot, & Villa-Vialaneix, 2017). The objective was to establish theoretical distributions of G’’st and φ’st allowing future predictions of values according to both *T* and *He*. For each mutation rate *µ* and each metric of genetic differentiation *GD* (either G’’st or φ’st) computed after the creation of the barrier (i.e., for T > 0), we used the R-package *randomForest* (Liaw & Wiener, 2002) to fit the model We used 200 trees and a sample size of 500, as these values provided very good accuracy (mean squared errors lower than 0.4%). Created *randomForest* R-objects were saved in the form of .*rda* files (the usual file format for saving R objects) and were further used to predict the four possible expected measures of genetic differentiation (two possible metrics of genetic differentiation and two possible mutation rates) between pairs of populations according to both the mean expected heterozygosity *He* (the proxy for effective population sizes) and the number of generations *T* elapsed since barrier creation, using the *predict*.*randomForest* function.

The 20460 simulations associated with T ≤ 0 (i.e., before the creation of the barrier) were used to predict the four possible measures of genetic differentiation (background signal) that may be expected under the influence of mutations, drift and incomplete genetic sampling between two adjacent populations not separated by any barrier to gene flow. For each of both mutation rates *µ* and each of both metrics of genetic differentiation *GD* (either G’’st or φ’st) computed before the creation of the barrier (i.e., for T < 0), was computed as the fifth percentile of non-null simulated *GD* values. These four predicted values were stored in the form of a .*rda* file.

### Computing the genetic index of fragmentation *F*_*INDEX*_

Equation 1 allows computing a unique index of fragmentation for each combination of both a mutation rate *µ* (5×10^−5^ or 5×10^−4^) and a metric of genetic differentiation *GD* (G’’st or φ’st). The four indices are then averaged to get the final index of fragmentation *F*_*INDEX*_ with a 95% confidence interval computed as, with the estimated standard error (i.e., the estimated standard deviation divided by 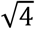).

Note that when several genotypic datasets are available for the same obstacle, for instance when several sympatric species are sampled on either side of the obstacle or when several replicates are considered (as is the case of all simulated data in this study), an overall *F*_*INDEX*_ can also be estimated using an intercept-only mixed-effect linear model with the various indices as the response variable and the genotypic dataset as a random effect (Bates, Mächler, Bolker, & Walker, 2015). This procedure allows taking into account the fact that *F*_*INDEX*_ values computed from the same dataset are not independent and thus avoids biased estimates of standard errors (McNeish, 2014). The overall *F*_*INDEX*_ is obtained from the estimated intercept of the model (which simply amounts to calculating the average of indices across datasets) and the corresponding 95% confidence interval is computed as, with the unbiased standard error as estimated from the mixed-effect model.

The whole procedure was automated within a user-friendly R-function (the FINDEX() R(- function; see Appendix S1). Users are simply expected to provide empirical genotypic datasets (in the *genepop* format) and a parameter file indicating for each considered obstacle the name of the two adjacent populations (as given in the genotypic datasets) and the number of generations elapsed since barrier creation. This number of generations is to be estimated from the life-history traits of the considered species. Figure 1 provides a flowchart allowing an overall visualization of the process.

**Figure 1.**
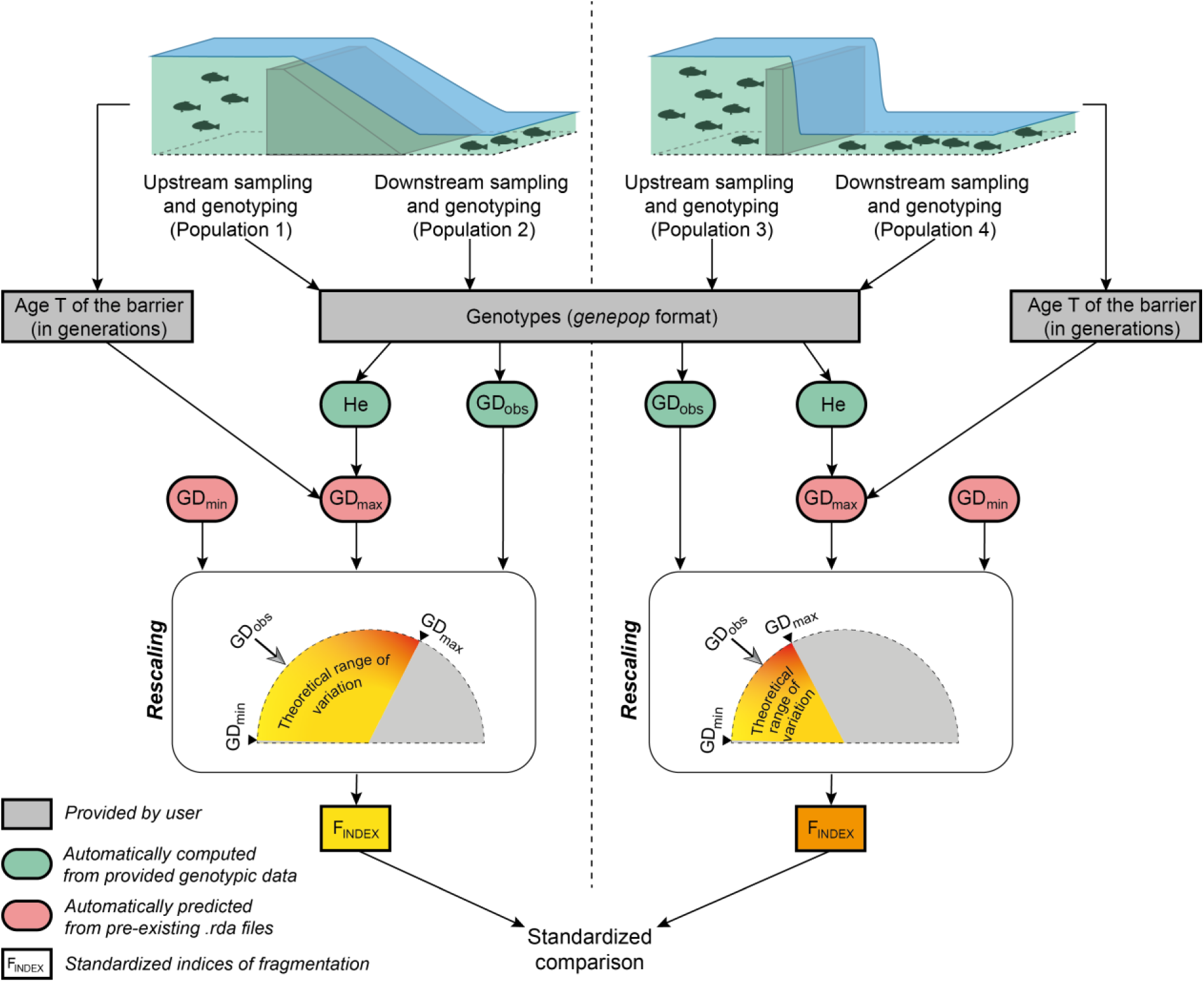
Flowchart illustrating the major steps in calculating the genetic index of fragmentation for two independent obstacles. This flowchart refers to a user-friendly script made publicly available. After the sampling of populations located at the immediate upstream and downstream vicinity of each obstacle, users only have to provide a file of genotypes in the *genepop* format and a file of parameters indicating, for each obstacle, the names of the sampled populations and the number *T* of generations elapsed since the creation of the obstacle. Observed measures of genetic differentiation and *GD*_*obs*_ mean expected heterozygosity *He* are automatically computed from provided genotypic data. *GD*_*min*_ and *GD*_*max*_ values, both delimiting the theoretical range of variation of *GD*_*obs*_, are automatically predicted from pre-existing .rda files, *GD*_*max*_ values depending on both *He* and *T*. The computation of the index basically amounts to rescaling *GD*_*obs*_ within its theoretical range (see main text for details), thus allowing standardized comparisons of the permeability of various obstacles, whatever their age, the considered species or the effective size of sampled populations.

### Validation of the *F*_*INDEX*_ from simulated data

To assess the validity of the proposed *F*_*INDEX*_ in response to different levels of obstacle permeability, we again used the program QuantiNemo2 to simulate gene flow over 1000 non-overlapping generations between two adjacent demes of constant carrying capacity *K*, with *K =* 50, 100, 250, 500 or 1000 individuals. To mimic realistic genetic datasets, each microsatellite locus was given a unique stepwise mutation rate *µ* randomly picked from a log-normal distribution ranging from 5×10^−5^ to 5×10^−3^ with a mean of 5×10^−4^ (see Appendix S7 for details). The inter-deme migration rate was set to 0.5 for the first 400 generations, and then dropped to *m* for the last 600 generations, with *m* ranging from 0 to 0.2 with an increment of 0.01 and from 0.2 to 0.5 with an increment of 0.05, mimicking the creation of a more or less severe barrier to gene flow (total barrier, crossing rate *m* = 0; no barrier, crossing rate *m* = 0.5). All other simulation parameters were similar to previous simulations. For each carrying capacity *K* and each crossing rate *m*, we ran 20 simulation replicates, and 30 genotypes were sampled at generations *t* = 405 (age of the barrier *T* = 5), 410, 415, 420, 425, 450, 500 and 700 (*T* = 300), resulting in a total of 21600 simulated genetic datasets in the *Fstat* format, further converted into the *genepop* format.

For each simulated dataset, we computed the averaged expected heterozygosity *He* and the two pairwise measures of genetic differentiation G’’st and φ’st. We then used parameters *T* and *He* to predict the corresponding measures of genetic differentiation and (for both G’’st and φ’st) expected under the two mutation rates 5×10^−5^ and 5×10^−4^ using the *predict*.*randomForest* function and the previously created .*rda* files (Appendix S1). For each dataset, the four indices of fragmentation were then computed using equation 1. To average datasets across replicates, we finally used intercept-only mixed-effect models (with dataset as a random effect) to get the final mean *F*_*INDEX*_ (along with a 95% confidence interval) corresponding to each combination of *K, T* and *m*.

We finally explored the sensitivity of the *F*_*INDEX*_ to uncertainty in the estimates of *Ne* and *T*. Details are provided in Appendices S12 and S13.

### Test of the *F*_*INDEX*_ with empirical data

To assess the behavior of the proposed *F*_*INDEX*_ in real situations, we used two published empirical datasets. The first one is from Gouskov et al. (2016). In this study, authors assessed riverscape fragmentation induced by 37 hydroelectric recent power stations in the Rhine catchment using data from 2133 European chubs (*Squalius cephalus*) sampled across 47 sites and genotyped at nine microsatellite loci. We selected 6 pairs of populations according to the following criteria: upstream and downstream populations belonged to the same river, were separated by a single dam, were distant from a maximum of 5km (the maximum migration distance observed in chubs being 16 km according to Fredrich et al., 2003) and were not separated by any confluence with important tributaries. This selection corresponded to 6 independent dams created between 1893 and 1964 (∼4 to 10 meters high), all equipped with a fishpass (Table1; see also Appendix S8 for a map). We considered a generation time of 3 years, as reported in Gouskov et al. (2016) to compute the number of generations elapsed since barrier creation and ran the developed FINDEX() R-function (Appendix S1) to automatically compute *F*_*INDEX*_ values.

The second empirical dataset is from Prunier et al. (2018). In this study, authors assessed the influence of various anthropogenic stressors including riverscape fragmentation induced by weirs on patterns of genetic diversity and differentiation in two freshwater fishes from two distinct rivers in southwestern France. They used data from 1361 Eurasian minnows (*Phoxinus phoxinus*) and 1359 Languedoc gudgeon (*Gobio occitaniae*) sampled across 47 sites (22 in the Célé River and 25 in the Viaur River) and genotyped at 11 and 13 microsatellite loci, respectively. We selected 8 pairs of populations according to the following criteria: upstream and downstream populations belonged to the same river, were separated by a single weir, were distant from a maximum of 1km, were not separated by any confluence with tributaries and were sampled for both species. This selection corresponded to 8 independent weirs (∼1 to 3 meters high) created between the 16th and the 20th century (Table 1; see also Appendix S8 for maps). We considered a generation time of 2 years in *P. phoxinus* and 2.5 years in *G. occitaniae* to compute the number of generations elapsed since barrier creation and again ran the FINDEX() R-function (Appendix S1) to automatically compute *F*_*INDEX*_ values for each obstacle, each species and across species.

**Table 1.** Main characteristics and results for the obstacles selected from empirical datasets (Original publication: (1) Gouskov et al., 2016; (2) Prunier et al., 2018). For each obstacle, the table indicates the name of the river, the date of creation, the distance between upstream and downstream sampled populations, the considered species (*Sc*: *Squalius cephalus*; *Go* : *Gobio occitaniae*; *Pp*: *Phoxinus phoxinus*), the number of generations elapsed since barrier creation, the mean expected heterozygosity (*He*), and the computed *F*_*INDEX*_ along with its 95% confidence interval. In bold, obstacles that were found as significant barriers to gene flow.

| River | Obstacle | Creation date | Upstream-Downstream distance (km) | Species | Number of elapsed generations | He | $F_{INDEX}$ | 95%CI | Original publication |
| --- | --- | --- | --- | --- | --- | --- | --- | --- | --- |
| **Rhine** | **Barr11** | **1964** | **4.79** | *Sc* | **15.33** | **0.69** | **55.22** | **5.71** | **(1)** |
| **Aar** | **Barr13** | **1902** | **1.91** | *Sc* | **36.00** | **0.76** | **49.53** | **10.72** | **(1)** |
| Aar | Barr17 | 1893 | 3.14 | *Sc* | 39.00 | 0.77 | 0 | 0 | (1) |
| **Aar** | **Barr19** | **1896** | **1.93** | *Sc* | **38.00** | **0.76** | **61.87** | **3.16** | **(1)** |
| Aar | Barr26 | 1963 | 4.84 | *Sc* | 15.67 | 0.75 | 0 | 0 | (1) |
| Limmat | Barr33 | 1933 | 3.24 | *Sc* | 25.67 | 0.72 | 0 | 0 | (1) |
| **Célé** | **CLA** | **1500** | **0.18** | *Go* | **204** | **0.60** | **42.53** | **3.88** | **(2)** |
| **Célé** | **SCA** | **1500** | **0.09** | *Go* | **204** | **0.63** | **35.09** | **2.60** | **(2)** |
| **Célé** | **SCC** | **1960** | **0.2** | *Go* | **20** | **0.64** | **64.97** | **2.81** | **(2)** |
| Viaur | SEG | 1600 | 0.11 | *Go* | 164 | 0.58 | 6.90 | 4.33 | (2) |
| Viaur | CAM | 1600 | 0.49 | *Go* | 164 | 0.62 | 0 | 0 | (2) |
| **Viaur** | **CAP** | **1700** | **0.55** | *Go* | **124** | **0.61** | **42.22** | **0.97** | **(2)** |
| **Viaur** | **SJU** | **1800** | **1.07** | *Go* | **64** | **0.62** | **45.01** | **3.16** | **(2)** |
| **Viaur** | **CIR** | **1960** | **0.97** | *Go* | **20** | **0.62** | **37.55** | **15.60** | **(2)** |
| **Célé** | **CLA** | **1500** | **0.18** | *Pp* | **255** | **0.54** | **36.12** | **3.80** | **(2)** |
| Célé | SCA | 1500 | 0.09 | *Pp* | 255 | 0.57 | 0 | 0 | (2) |
| Célé | SCC | 1960 | 0.2 | *Pp* | 25 | 0.58 | 10.05 | 11.41 | (2) |
| Viaur | SEG | 1600 | 0.11 | *Pp* | 205 | 0.63 | 12.03 | 13.61 | (2) |
| Viaur | CAM | 1600 | 0.49 | *Pp* | 205 | 0.61 | 0 | 0 | (2) |
| Viaur | CAP | 1700 | 0.55 | *Pp* | 155 | 0.67 | 0 | 0 | (2) |
| Viaur | SJU | 1800 | 1.07 | *Pp* | 105 | 0.70 | 0 | 0 | (2) |
| Viaur | CIR | 1960 | 0.97 | *Pp* | 25 | 0.70 | 0 | 0 | (2) |
| **Célé** | **CLA** | **1500** | **0.18** | ***Go-Pp*** | **/** | **/** | **39.32** | **6.23** | **(2)** |
| Célé | SCA | 1500 | 0.09 | *Go-Pp* | / | / | 17.55 | 34.39 | (2) |
| Célé | SCC | 1960 | 0.2 | *Go-Pp* | / | / | 37.51 | 53.83 | (2) |
| Viaur | SEG | 1600 | 0.11 | *Go-Pp* | / | / | 9.47 | 6.88 | (2) |
| Viaur | CAM | 1600 | 0.49 | *Go-Pp* | / | / | 0 | 0 | (2) |
| Viaur | CAP | 1700 | 0.55 | *Go-Pp* | / | / | 21.11 | 41.38 | (2) |
| Viaur | SJU | 1800 | 1.07 | *Go-Pp* | / | / | 22.51 | 44.12 | (2) |
| Viaur | CIR | 1960 | 0.97 | *Go-Pp* | / | / | 18.78 | 36.80 | (2) |

## Results

### Expected measures of genetic differentiation

The first set of simulations was designed to predict *GD*_*min*_ and *GD*_*max*_ values, that is, the lower and upper limits of the theoretical range of variation of *GD*_*obs*_. Data simulated before the creation of the barrier (*m* = 0.5; *t* < 400; T < 0) were used to predict *GD*_*min*_ values whereas data simulated after the creation of the barrier (*m* = 0; *t* ≥ 400; T ≥ 0) were used to predict *GD*_*max*_ values. As expected with a migration rate of 0.5, *GD*_*min*_ values were always very close from 0 (∼0.8 10^−3^ for G’’st, ∼1.2 10^−3^ for φ’st; Appendix S6). These values represent the predicted background levels of genetic differentiation resulting from the sole influences of random processes such as genetic drift, mutations and sampling biases (Figure 2).

**Figure 2.**
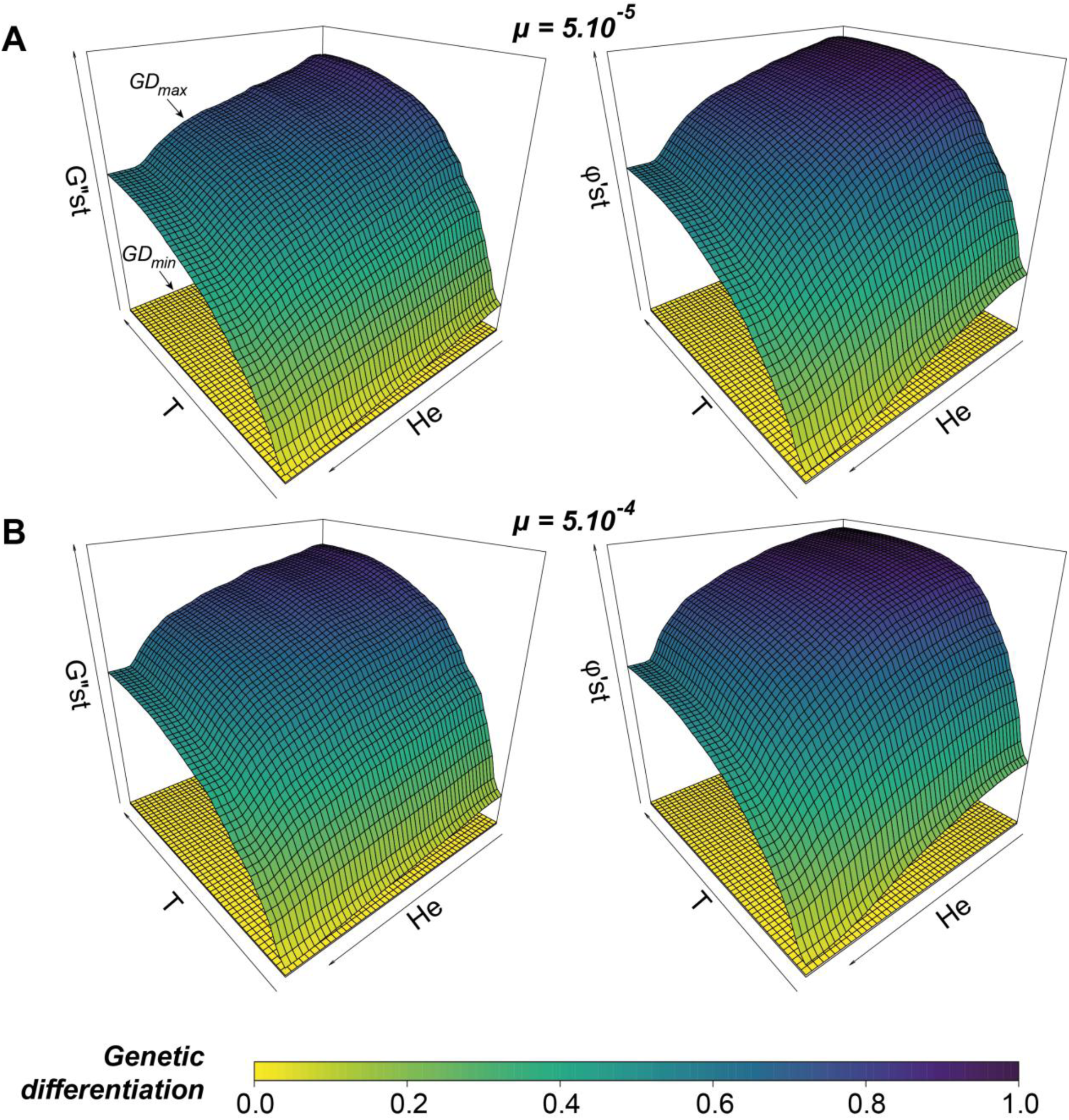
For each mutation rate (panels A and B) and each metric of genetic differentiation (G’’st on the left and φ’st on the right), predicted *GD*_*max*_ variations across the parameter space defined by the number *T* of generations elapsed since total barrier creation (from 0 to 600 generations) and the averaged expected heterozygosity (*He*, ranging from 0 to 0.93) for pairs of adjacent populations. *GD*_*min*_ values are represented at the bottom of each graph. *GD*_*min*_ and *GD*_*max*_ surfaces together delimit the theoretical range of variation for any observed measure of genetic differentiation *GD*_*obs*_.

*GD*_*max*_ values were on the contrary designed to mimic the temporal inertia in the setting up of genetic differentiation after the creation of a total barrier to gene flow. They were predicted from the number T of elapsed generations since barrier creation and the averaged expected heterozygosity *He* from simulated data using a Random Forest algorithm. With explained variance ranging from 86.8 to 94.2 %, Random Forest models accurately captured variations in measures of genetic differentiation across the parameter space, whatever the considered mutation rate or the considered metric of genetic differentiation (see Appendices S9 and S10). As expected in absence of gene flow (Figure 2), *GD*_*max*_ increased with time since barrier creation and decreased with the increase in effective population size (i.e., with *He*). With predicted *GD*_*max*_ values ranging from 0.031 to 0.898 for G’’st and from 0.042 to 0.968 for φ’st, both metrics displayed similar distribution patterns across mutation rates, although φ’st systematically showed higher values at low *He*.

### Validation of the *F*_*INDEX*_ from simulated data

The second set of simulations was designed to assess whether the *F*_*INDEX*_ correctly reflected the actual level of gene flow between two populations separated by an artificial barrier, beyond the temporal inertia in the setting-up of genetic differentiation. The mean *F*_*INDEX*_ values computed over simulated replicates for each combination of *K* (carrying capacity), *T* (number of generations since barrier creation) and *m* (obstacle crossing rate) showed –as expected– an overall decrease with the increase in crossing rate, whatever the size of populations or the age of the barrier (Figure 3A-D). As expected when population are connected with high crossing rates (*m* > 0.2, a crossing rate of 0.5 leading to full connectivity), the 95% confidence intervals about the *F*_*INDEX*_ always included values lower than 20%. On the contrary, in absence of gene flow (*m* = 0), the 95% confidence intervals always included values higher than 90%, except within the first 10 generations after barrier creation (Figure 3A-B). In these cases, the *F*_*INDEX*_ was slightly biased downwards, which indicates that we could not totally rule out the noise associated with the measurement of genetic differentiation within the 10 first generations after barrier creation (Appendix S11). Nevertheless, the *F*_*INDEX*_ showed valid and consistent values for both lowest and highest crossing rates, the two thresholds of 90% (total barrier to gene flow) and 20% (full gene flow) providing robust benchmarks for future interpretation of the index, whatever the age of the obstacle (from generation 10 at least) or the effective size of populations.

**Figure 3.**
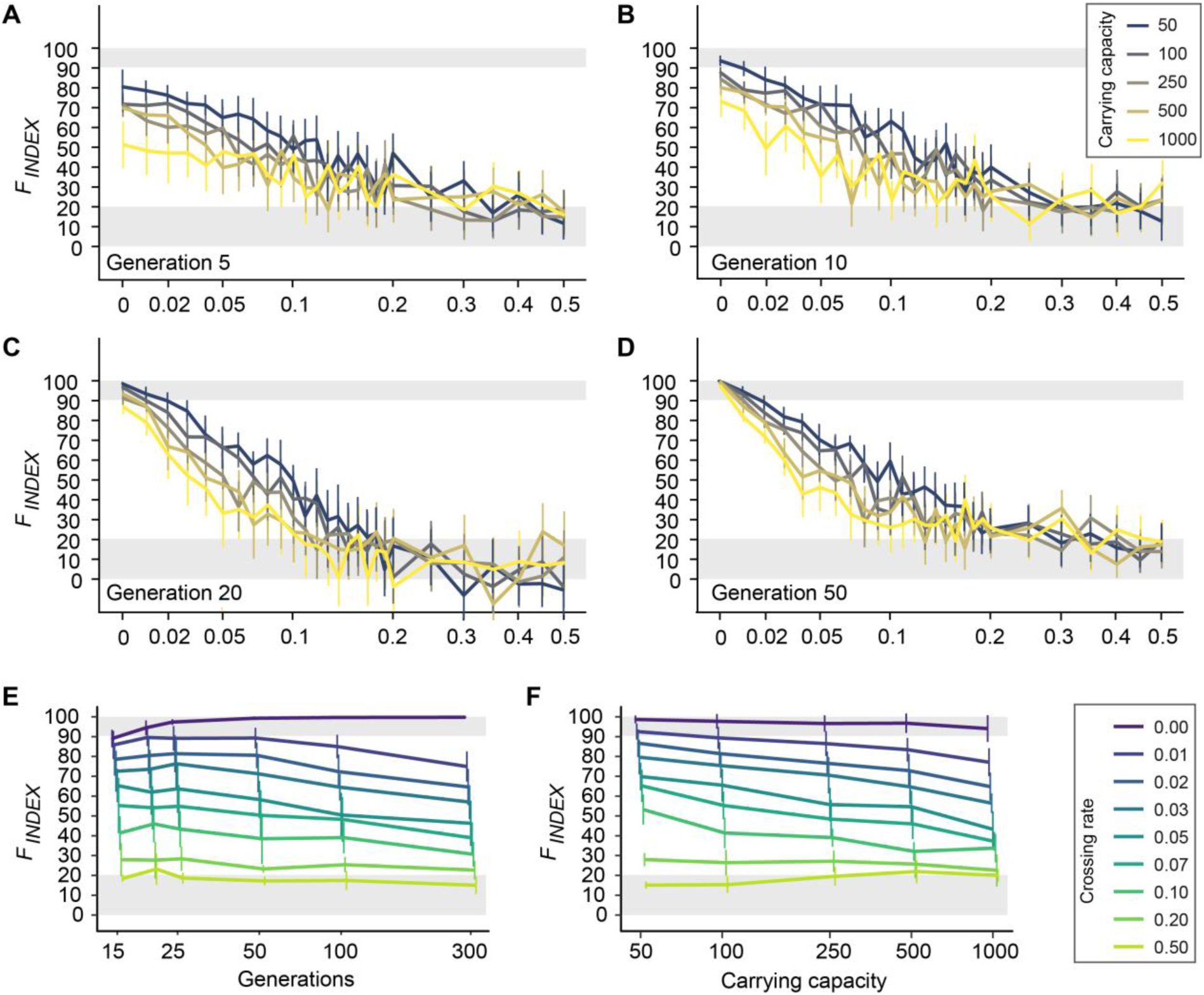
Panels A-D: *F*_*INDEX*_ responses to the increase in crossing rate (*m*, on a logarithmic scale) for five different carrying capacities (colored lines) and from 5 to 50 generations after barrier creation. Results for a number of generations higher than 50 are visually similar to panel D (not shown; but see panel E). All *F*_*INDEX*_ values were averaged over 20 simulated replicates and plotted with 95% confidence intervals. Panels E-F: *F*_*INDEX*_ responses to the increase in time since barrier creation (panel E) and to the increase in carrying capacity (panel F) for eight different crossing rates *m* (colored lines). The mean *F*_*INDEX*_ values computed over simulated replicates were here averaged over carrying capacities (panel E) or over generations (excluding generations ≤ 10; panel F) and plotted with standard deviations. In all panels, shaded grey areas represent the ranges of variations in which the monitored obstacle can be considered as acting as a total barrier to gene flow (*F*_*INDEX*_ > 90%) or, on the contrary, as allowing full genetic connectivity (*F*_*INDEX*_ < 20%).

For low –though non-null– crossing rates (0 < *m ≤* 0.1), the *F*_*INDEX*_ showed higher variability, with two noticeable trends. First, whatever the simulated carrying capacity, the *F*_*INDEX*_ showed a slight 10 to 20% decrease with the increase in time since barrier creation (from generation 15 to generation 300; Figure 3E). For a crossing rate of *m* = 0.05 for instance, *F*_*INDEX*_ values decreased from 65% at generation 15 to ∼46% at generation 300. Secondly, whatever the generation (> 10), the *F*_*INDEX*_ showed a slight 10 to 30% decrease with the increase in effective population sizes (from carrying capacity K = 50 to 1000; Figure 3F). For a crossing rate of *m* = 0.05 for instance, *F*_*INDEX*_ values decreased from 70% in smallest populations (K = 50) to ∼43% in largest populations (K=1000).

Sensitivity analyses finally showed that the *F*_*INDEX*_ is highly robust to a ∼50% uncertainty in the estimates of *T* and that 95% CI about *F*_*INDEX*_ values correctly capture uncertainty associated with the use of *He* as a proxy for *Ne*. Details are provided in Appendices S12 and S13.

### Test of the *F*_*INDEX*_ with empirical data

In the first empirical dataset (Gouskov et al., 2016), monitored dams were created from 1893 to 1964, which corresponds to approximately 15 to 39 generations in *S. cephalus* (Table1). Averaged levels of expected heterozygosity were high and showed little variability (ranging from 0.69 to 0.77), whereas observed measures of genetic differentiation were pretty low, ranging from 0 to 0.028 for φ’st and from 0 to 0.025 for G’’st. We found that three dams showed a *F*_*INDEX*_ value ranging from 49% to 62%, suggesting a 49 to 62% local decrease in genetic connectivity (Figure 4A). The other three dams all showed null *F*_*INDEX*_ values, indicating that populations located on either side of the barrier are fully connected by gene flow (Table 1). Importantly, *F*_*INDEX*_ values were independent from both time since barrier creation (spearman correlation test, *ρ* = 0.03, p = 0.95) and averaged expected heterozygosity (*ρ* = −The first set of simulations was designed to predic0t.03, p = 0.95).

**Figure 4.**
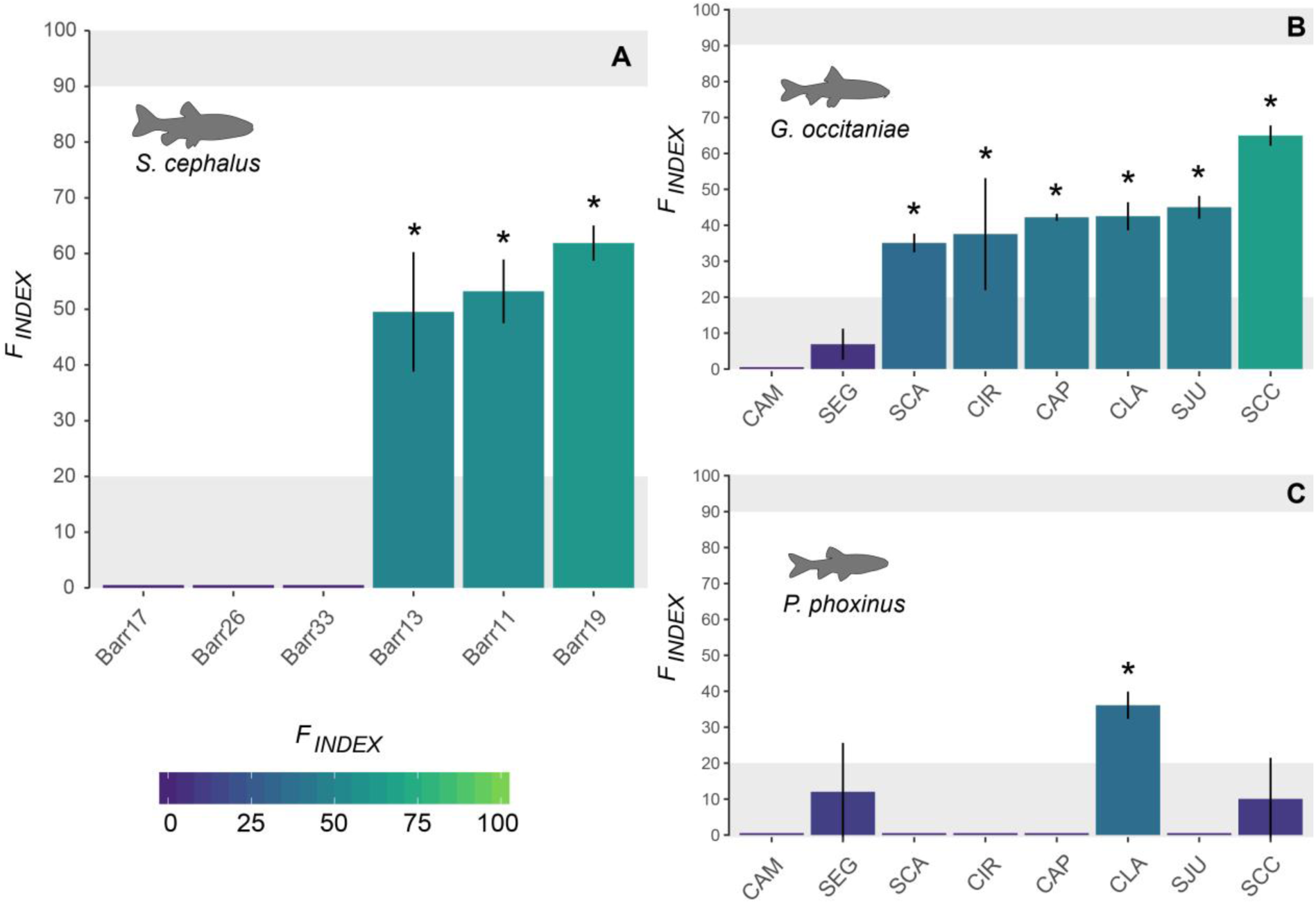
*F*_*INDEX*_ values and associated 95% confidence intervals as computed from empirical genetic datasets in chubs (panel A), gudgeons (panel B) and minnows (panel C). In all panels, shaded grey areas represent the ranges of variations in which the monitored obstacle can be considered as acting as a total barrier to gene flow (*F*_*INDEX*_ > 90%) or, on the contrary, as allowing full genetic connectivity (*F*_*INDEX*_ < 20%). Stars indicate a significant barrier effect of the obstacle. See Table 1 for details.

In the second empirical dataset (Prunier et al., 2018), monitored weirs were built between the 16th and the 20th century, that is approximately from 20 to 204 generations in *G. occitaniae* and from 25 to 255 generations in *P. phoxinus*. As previously, averaged levels of expected heterozygosity were high and showed little variability (ranging from 0.58 to 0.72), whereas observed measures of genetic differentiation were pretty low, ranging from 0 to 0.034 for φ’st and from 0 to 0.026 for G’’st. The impact of weirs was variable across space and species (Table 1; Figure 4B). In *G. occitaniae*, six weirs (out of eight) were found as responsible for a decrease in genetic connectivity since barrier creation (*F*_*INDEX*_ > 20%), with *F*_*INDEX*_ values ranging from 35% in the case of barrier SCA to 65% in the case of barrier SCC in the Célé River. In *P. phoxinus*, all weirs but CLA in the Célé River (*F*_*INDEX*_ = 36%) were found as highly permeable (*F*_*INDEX*_ < 20%), with five out of eight weirs showing a *F*_*INDEX*_ of 0%. When computed across species, only the barrier CLA in the Célé River (multispecies *F*_*INDEX*_ = 39%) was identified as an obstacle to overall genetic connectivity (other *F*_*INDEX*_ values ranging from 0 to %, with 95% confidence intervals systematically including the 20% threshold; Table 1). As previously, *F*_*INDEX*_ values in each species were independent from both time since barrier creation (|*ρ*| < 0.46, p > 0.25) and averaged expected heterozygosity (|*ρ*| < 0.54, p > 0.16).

## Discussion

Restoring riverscape connectivity is of crucial importance in terms of biodiversity conservation and it is now often subject to regulatory obligations (e.g. in Europe, the Water Framework Directive 2000/60/EC). However, rivers are subject to many and sometimes contradictory uses (Reid et al., 2018): for practitioners to be able to propose informed trade-offs between restoring riverscape connectivity and maintaining infrastructures and their associated socioeconomic benefits (Hand et al., 2018; Roy et al., 2018; Song et al., 2019), new tools have to be developed, allowing a rapid and reliable quantification of the relative impacts of obstacles to freshwater species movements. Our objective was here develop an operational tool allowing such thorough quantification from a minimum amount of fieldwork and data (Figure 1; see Box 1 for user guidelines).

#### Box 1

**Guidelines for the use and the interpretation of the *F*_*INDEX*_.**

The *F*_*INDEX*_ allows an individual and standardized quantification of the impact of artificial barriers on riverscape functional connectivity from snapshot measures of genetic differentiation. Here, we provide a guideline for practitioners:

- **Species**: any freshwater species whose local effective population sizes are lower than 1000 can be considered.
- **Obstacle:** any obstacle whose age corresponds to a minimum of 10-15 generations and a maximum of 600 generations for the studied species can be considered.
- **Sampling:** populations are sampled in the immediate upstream and downstream vicinity of the obstacle, with a minimum of 20-30 individuals per populations.
- **Genetic data**: Individual genotypes are based on a set of polymorphic microsatellite markers.
- **Computation:** The *F*_*INDEX*_ is computed in R thanks to a user-friendly script made publicly available (see Data Archiving statement and Appendix S1 for a walkthrough).
- **Interpretation for** *F*_*INDEX*_ **> 90%:** A *F*_*INDEX*_ value higher than 90% (or whose 95% confidence interval includes the 90% threshold) indicates no gene flow between populations (total barrier effect), whatever the age of the obstacle or the effective size of populations.
- **Interpretation for** *F*_*INDEX*_ **< 20%:** A *F*_*INDEX*_ value lower than 20% (or whose 95% confidence interval includes the 20% threshold) indicates full genetic connectivity (no barrier effect), whatever the age of the obstacle or the effective size of populations.
- **Interpretation for intermediate** *F*_*INDEX*_ **values:** Intermediate *F*_*INDEX*_ values can be used to rank obstacles according to their barrier effect. However, for *F*_*INDEX*_ values ranging from ∼40 to ∼80%, the *F*_*INDEX*_ tends to slightly decrease with both the increase in the number of generations since barrier creation and the increase in effective population sizes (Figure 3E-F). Obstacles with *F*_*INDEX*_ values that do not differ by more than 15 to 20% but that are characterized by very different ages and / or population sizes (as indicated for instance by large differences in expected heterozygosity) should be considered as possibly having comparable barrier effects, except of course if the ranking of obstacles based on *F*_*INDEX*_ values goes against these trends. Consider for instance an obstacle A of age 20 (in generations) and an obstacle B of age 300. If *F*_*INDEX*_(A) = 40% and *F*_*INDEX*_(B) = 20%, both obstacles should be considered as possibly having the same impact on gene flow. On the contrary, if *F*_*INDEX*_(A) = 20% and *F*_*INDEX*_(B) = 40%, obstacle B can be confidently considered as more impactful than obstacle A. Similarly, consider an obstacle C separating populations with low expected heterozygosity (suggesting small effective population size) and an obstacle D separating populations with high expected heterozygosity. If *F*_*INDEX*_(C) = 40% and *F*_*INDEX*_(D) = 20%, both obstacles should be considered as possibly having the same impact on gene flow. On the contrary, if *F*_*INDEX*_(C) = 20% and *F*_*INDEX*_(D) = 40%, obstacle D can be confidently considered as more impactful than obstacle C.

The proposed genetic index of fragmentation *F*_*INDEX*_ can be easily and automatically computed from a simple set of upstream and downstream genotypes collected once and in the direct vicinity of a putative barrier, provided the approximate number of generations elapsed since barrier creation is known (the *F*_*INDEX*_ being highly robust to uncertainty in time since barrier creation; Appendix S13). Based on two complementary metrics of genetic differentiation (G’’st and φ’st) preliminary chosen so as to limit any possible bias, the *F*_*INDEX*_ simply scales the observed level of genetic differentiation (*GD*_*obs*_) with respect to a theoretical range of variation spanning from the background noise expected in the absence of any barrier to gene flow (*GD*_*min*_ ∼ *F*_*INDEX*_ = 0%) to the maximal level of differentiation expected if the obstacle was a total barrier to gene flow (*GD*_*max*_, *F*_*INDEX*_ = 100%). The latter takes into account both the time since barrier creation and the effective size of populations, which makes the *F*_*INDEX*_ a truly innovative tool as it makes it possible to compare the actual barrier effect of obstacles differing by their age and/or by the size of the populations they separate. Using numerous simulations, we were able to obtain *GD*_*max*_ values for a large range of biologically realistic parameters (Figure 2). As expected, *GD*_*max*_ values progressively increased with time since barrier creation and decreased with the increase in averaged expected heterozygosity. Mutation rate also influenced *GD*_*max*_ patterns: as expected, higher mutation rates accelerate genetic differentiation through time when population sizes are small to medium. The use of two realistic mutation rates in *GD*_*max*_ predictions allows taking into consideration uncertainty in our proxy for effective population sizes, in the form of a 95% confidence interval about *F*_*INDEX*_ values (see Appendice S5b and S12).

The *F*_*INDEX*_ showed constant patterns of decrease with the increase in crossing rates (from *m* = 0 to *m* = 0.2), whatever the number of generations since barrier creation and the effective population size (Figure 3). For the lowest crossing rates (m ≤ 0.05), we found however that it could underestimate barrier effects in the first 5 to 10 generations after the creation of the obstacle. As a conservative strategy, we suggest that the *F*_*INDEX*_ should not be used to assess the permeability of obstacles separating populations for fewer than 10 generations. However, it is noteworthy that the *F*_*INDEX*_ can be applied to any type of organisms and thus that species with short generation time (such as some invertebrate species) may be considered as good candidates to investigate the impact of recently built barriers (e.g., less than 10 years ago). For the lowest crossing rates (m ≤ 0.1), we also found that *F*_*INDEX*_ values slightly decreased with both time since barrier creation (from generations 15 to 300) and effective population sizes (from simulated carrying capacities K = 50 to K = 1000; Figure 3E-F). These trends have to be kept in mind when comparing intermediate *F*_*INDEX*_ values ranging from ∼40 to ∼80% (see Box 1 for guidelines).

Nevertheless, the *F*_*INDEX*_ provides a promising individual quantification of both the short- and long-term genetic effects of dam-induced fragmentation, allowing robust comparisons among species or populations with different population sizes, and obstacles of different ages (from generation 15 at least) and types. When applied to empirical data, the *F*_*INDEX*_ allowed identifying several obstacles partially limiting gene flow in the three considered freshwater fish species (Figure 4). In each dataset, computed *F*_*INDEX*_ values were systematically independent from both time since barrier creation and averaged expected heterozygosity, indicating that the *F*_*INDEX*_ properly takes into account the differential evolution of allelic frequencies on either side of the barrier. Interestingly, the SCC weir on the Célé River (Prunier et al., 2018) showed contrasting results in gudgeons and minnows: it was identified as the most impactful obstacle in gudgeons (*F*_*INDEX*_ = 65%; Table 1) whereas it was found as highly permeable to gene flow in minnows (*F*_*INDEX*_ = 10%). More generally, minnows were much less affected by obstacles than gudgeons, in accordance with personal field observations and previous findings on the same two rivers (but from independent datasets; Blanchet et al., 2010). Although understanding how obstacle typological features (height, slope, presence of a secondary channel, etc.; Baudoin et al., 2014) and fish traits (body size, movement capacities, etc.; Blanchet et al., 2010) might interact and shape riverscape patterns of functional connectivity was beyond the scope of this study, these results suggest that future comparative studies based on the proposed *F*_*INDEX*_ might provide thorough insights as to the determinants of dam-induced fragmentation in various freshwater organisms (Richardson, Brady, Wang, & Spear, 2016), including fish but also other taxa such as macro-invertebrates that display very contrasting traits related to dispersal (e.g., Alp, Keller, Westram, & Robinson, 2012).

Despite its strong operational potential, the *F*_*INDEX*_ however does not come without some limitations (see Box 2 for a list of possible future developments). First of all, it is important to remember that this index is a measure of genetic connectivity, not demographic connectivity (Lowe & Allendorf, 2010), and thus cannot directly provide any counting of the actual number of crossing events. If immigrants do not reproduce, the actual crossing of dozens of individuals, although suggesting high permeability, might not translate into low *F*_*INDEX*_ values. Although this is more of an inherent characteristic of the index than a real limitation, it is important to keep this specificity in mind when interpreting it. Furthermore, a crossing rate has to be interpreted in regard of effective population sizes: a crossing rate of 0.05 actually corresponds to 2.5 effective dispersal events per generation in populations of size 50, but to 50 effective dispersal events in populations of size 1000. This higher permeability in the latter case translates into *F*_*INDEX*_ values being systematically slightly lower when simulated population sizes are larger (at a given intermediate crossing rate; Figure 3F). However, since the actual effective size of natural populations is generally unknown, these differences in *F*_*INDEX*_ values may be difficult to interpret when handling empirical data. We provide guidelines for the interpretation of *F*_*INDEX*_ values in Box 1.

#### Box 2

**Future directions for improving the *F*_*INDEX*_.**

The *F*_*INDEX*_ is already operational but it is however still in its infancy. We identified several research avenues that may allow further improving it or help answer specific needs. They are here presented by our order of priority.

- **Taking asymmetric crossing into consideration:** The proposed *F*_*INDEX*_ currently relies on the use of classical pairwise measures of genetic differentiation that assume symmetric gene flow. It will be first necessary to assess the sensitivity of the current version of the *F*_*INDEX*_ to asymmetric barrier effects and, if needed, to determine whether existing asymmetric measures of genetic differentiation (e.g., Sundqvist, Keenan, Zackrisson, Prodöhl, & Kleinhans, 2016) could be used to improve its efficiency. This task may otherwise require the development of new metrics of genetic differentiation.
- **Dealing with non-adjacent sampling designs:** The proposed *F*_*INDEX*_ relies on a strict *adjacent sampling strategy*, with populations sampled in the immediate upstream and downstream vicinity of the considered obstacle. However, this sampling design might be difficult to implement in some situations (e.g., dams with a large reservoir). When the two sampled populations are distant from each other, values may nevertheless result from the interplay between the actual barrier effect (the quantity of interest) and other processes such as Isolation-by-Distance. In such situations, the *F*_*INDEX*_ should be computed using *ad hoc* and values, both taking into account the additional processes responsible for values. To that aim, a solution could be to consider a *space-for-time substitution sampling design* (e.g., Coleman et al., 2018), with the additional sampling of (at least) two control populations that are not disconnected by any barrier, are located within the same river stretch and are separated by approximately the same distance as the two focal populations. Measures of genetic differentiation computed between these control populations could be directly used as new *ad hoc* values. An empirical migration rate *m* could then be inferred from these control measures of genetic differentiation, for instance using an Approximate Bayesian Computation approach (Bertorelle, Benazzo, & Mona, 2010; Csilléry, Blum, Gaggiotti, & François, 2010) and new genetic simulations be performed using *m* as the baseline migration rate, so as to get *ad hoc* values (taking into consideration the age of the obstacle and the effective size of focal populations). Such a procedure might help answer very specific needs, but its complexity might restrict the direct use of the *F*_*INDEX*_ to informed and trained managers. Furthermore, additional work would be required to determine to what extent *F*_*INDEX*_ values computed in that way could still be comparable across obstacles.
- **Handling other genetic markers:** The proposed *F*_*INDEX*_ relies on the use of microsatellite markers. Microsatellite markers are still widely used in the study of non-human organisms, especially by environmental managers. However, the reduction in sequencing costs and the development of an ever-increasing supply of biotechnological services (Davey et al., 2011) now allow easier access to new genetic markers such as SNPs. In the future, a new version of the *F*_*INDEX*_ allowing the use of SNPs could be developed: it would require identifying relevant SNP-based measures of genetic differentiation (see Appendices S3 and S4) as well as numerous simulations to establish theoretical distributions of values according to both time since barrier creation and expected heterozygosity.

Secondly, the computation of the *F*_*INDEX*_ relies on the assumption that, beyond the background signal of genetic differentiation that is expected under the sole influences of genetic drift, mutations and incomplete genetic sampling (*GD*_*min*_), the observed measures of genetic differentiation only stem from dam-induced fragmentation. In other words, it is crucial to consider situations in which the focal populations would be fully connected if the obstacle did not exist. This assumption is true only when sampled populations are adjacent, that is, located in the immediate upstream and downstream vicinity of the obstacle (Figure 1). Although restrictive, this adjacent sampling design has the advantage of making the *F*_*INDEX*_ valid for almost any freshwater species, regardless of their life history traits: the effective population size, a key-parameter that may obviously differ across species, is indeed directly taken into consideration in the *F*_*INDEX*_ computation, while differences in dispersal abilities can be considered as null at very short distances. It yet implies the exclusion of migratory fish species, though at the heart of great conservation issues (e.g., Junge et al., 2014; Klütsch et al., 2019): complex life-cycles such as anadromy (“river-sea-river” migrations), catadromy (“sea-river-sea” migrations) or potamodromy (“river-lake-river” migrations) indeed preclude the delineation of upstream and downstream populations and do not allow proper estimates for the *F*_*INDEX*_. In non-migratory fish species, this assumption also prevents the use of the *F*_*INDEX*_ in large-scale studies, in which the distance between the upstream and the downstream sampling sites lies beyond the dispersal capacities of the studied species. It certainly leaves room for manoeuvre, as illustrated with the empirical dataset from Gouskov et al. (2016): we could for instance select pairs of populations located up to 5km apart, but this was only possible because of the high mobility of chubs, and performed in an illustrative purpose: a maximum distance of 1km would have been safer. In low-mobility species, a non-adjacent sampling design might bias the *F*_*INDEX*_ upwards and hence overestimate the effect of obstacles, as observed measures of genetic differentiation would result from dam-induced fragmentation but also from other processes such as Isolation-by-Distance (e.g. Coleman et al., 2018). We thus strongly encourage practitioners to consider an adjacent sampling design as often as possible, although we readily acknowledge that this may not always be an easy task given safety and accessibility considerations. Furthermore, fish may not always be found in the direct vicinity of obstacles. For instance, the conversion of a river into a reservoir after the creation of a dam often leads to major habitat modification and shifts in species composition (Bednarek, 2001), which can force adapting the sampling design. A solution might be to capture the resultant background signal of genetic differentiation by simulating *ad hoc GD*_*min*_ and *GD*_*max*_ values under various scenarios of isolation (Isolation-by-Distance, Isolation-by-Resistance, etc.; McRae, 2006) in a way similar to the simulation of *GD*_*max*_ values in this study (Figure 2; see also Box 2). It is in this perspective that the provided R-function already allows users to integrate their own *GD*_*min*_ and *GD*_*max*_ values (Appendix S1). However, we believe that the variety, the complexity and the specificity of such scenarios would preclude the computation of standardized *F*_*INDEX*_ scores, comparable among obstacles, species and studies. Although it might in some instances be considered a technical constraint, we argue that only a strict adjacent sampling design can warrant unbiased and reliable *F*_*INDEX*_ estimates.

Finally, the proposed *F*_*INDEX*_ does not take into account the possible asymmetric gene flow created by barriers, as fish might struggle or even fail to ascent an obstacle (sometimes despite the presence of dedicated fishpasses; Silva et al., 2018) whereas dam discharge might on the contrary further increase or even force downstream movements (Pracheil et al., 2015). Although quantifying the asymmetric permeability of obstacles appears of crucial importance for informed conservation measures, the proposed *F*_*INDEX*_ currently relies on the use of classical pairwise measures of genetic differentiation that assume symmetric gene flow. This may for instance partly explain why we did not find any *F*_*INDEX*_ higher than 65% for weirs (Prunier et al., 2018) and 61% for dams (Gouskov et al., 2016; Table 1), a result that calls for future comparisons of the *F*_*INDEX*_ with direct monitoring methods (Cayuela et al., 2018). Future developments will be required to allow the *F*_*INDEX*_ to provide unbiased and distinct standardized scores for both upstream and downstream barrier effects (see Box 2). In the meanwhile, it may be interesting to also assess the validity of the *F*_*INDEX*_ in quantifying the effects of terrestrial obstacles, since asymmetric gene flow is not necessarily as pronounced as in river systems: provided that populations are sampled in the direct vicinity of the obstacle, the *F*_*INDEX*_ might as well provide a standardized quantification of road-induced fragmentation.

## Conclusion

We here laid the groundwork for an operational tool dedicated to the individual and standardized quantification of the impact of artificial barriers on riverscape functional connectivity from measures of genetic differentiation. The proposed genetic index of fragmentation *F*_*INDEX*_ is designed to take into account the temporal inertia in the evolution of allelic frequencies resulting from the interplay between the age of the obstacle and the effective sizes of populations. Provided only adjacent populations are sampled, the *F*_*INDEX*_ allows a rapid and thorough ranking of obstacles only a few generations after their creation. The *F*_*INDEX*_ in its current form still suffers from some limitations, and it should be seen as the preliminary version of a future powerful bio-indicator of habitat fragmentation, rather than as an end-product. We call conservation and population geneticists to pursue the development of such an index, as we –as scientists– need to help managers resolve complex and urging social problems. In Box 2, we hence propose several research avenues. Nonetheless, the *F*_*INDEX*_ is robust, only requires a minimum amount of fieldwork and genotypic data and already solves several difficulties inherent to the study of dam-induced fragmentation in river systems, making it a promising tool for the restoration of riverscape connectivity. The *F*_*INDEX*_ may allow practitioners to objectively identify obstacles that do not present any substantial conservation issue (from a connectivity perspective) and help them target their efforts and resources towards the most impactful ones. Similarly, it may allow tracking the expected temporal decrease in genetic differentiation after obstacle removal or fishpass setting and thus help evaluate the success of local mitigations and restoration measures in response to regulatory obligations. Finally, it might as well provide a standardized quantification of road-induced fragmentation, a critical issue in terrestrial ecology.

## Acknowledgements

We warmly thank all the colleagues and students who helped with field sampling. We are also grateful to Dr. A. Gouskov and C. Vorburger for details about their data. This work has been financially supported by grants to SB from the Agence Française pour la Biodiversité and from the Région Occitanie (CONAQUAT).

## Data Archiving

The two simulated datasets as well as R-objects allowing the computation of the *F*_*INDEX*_ are available at the Figshare Digital Repository (https://doi.org/10.6084/m9.figshare.9698879.v4) as well as on our personal website (http://www.jeromeprunier.fr/Tools.html).

## Supporting Information

Appendix S1: The FINDEX() R-function: Walkthrough for the computation of the *F*_*INDEX*_

Appendix S2: Details on the *F*_*INDEX*_ formula.

Appendix S3: Effect of sample size on various measures of genetic differentiation.

Appendix S4: Effect of both mutation rate and time since barrier creation on various measures of genetic differentiation.

Appendix S5: Relationship between mean expected heterozygosity and mean effective population size.

Appendix S6: Estimates of values.

Appendix S7: Distribution of mutation rates used for validation of the index (second set of simulations).

Appendix S8: Localization of obstacles in the two empirical datasets.

Appendix S9: Relationship between observed and predicted measures of genetic differentiation.

Appendix S10: Variable importance in Random Forest predictive models.

Appendix S11: Rational for the observed bias in *F*_*INDEX*_ values in the very first generations after barrier creation.

Appendix S12: Robustness of the *F*_*INDEX*_ to uncertainty in effective population sizes.

Appendix S13: Robustness of the *F*_*INDEX*_ to uncertainty in the number of generations elapsed since barrier creation.

## Literature Cited

Allendorf, F. W. (1986). Genetic drift and the loss of alleles versus heterozygosity. Zoo Biology, 5, 181–190.

Alp, M., Keller, I., Westram, A. M., & Robinson, C. T. (2012). How river structure and biological traits influence gene flow: A population genetic study of two stream invertebrates with differing dispersal abilities: Biological traits and gene flow in stream invertebrates. Freshwater Biology, 57, 969–981.

Bates, D., Mächler, M., Bolker, B., & Walker, S. (2015). Fitting Linear Mixed-Effects Models Using lme4. Journal of Statistical Software, 67. https://doi.org/10.18637/jss.v067.i01

Baudoin, J.-M., Burgun, V., Chanseau, M., Larinier, M., Ovidio, M., Sremski, W., … Voegtle, B. (2014). The ICE protocol for ecological continuity. Assessing the passage of obstacles by fish. Concepts, design and application. Paris: Onema.

Bednarek, A. T. (2001). Undamming Rivers: A Review of the Ecological Impacts of Dam Removal. Environmental Management, 27, 803–814.

Bertorelle, G., Benazzo, A., & Mona, S. (2010). ABC as a flexible framework to estimate demography over space and time: Some cons, many pros. Molecular Ecology, 19, 2609–2625.

Birnie-Gauvin, K., Aarestrup, K., Riis, T. M. O., Jepsen, N., & Koed, A. (2017). Shining a light on the loss of rheophilic fish habitat in lowland rivers as a forgotten consequence of barriers, and its implications for management. Aquatic Conservation: Marine and Freshwater Ecosystems, 27, 1345–1349.

Blanchet, S., Rey, O., Etienne, R., Lek, S., & Loot, G. (2010). Species-specific responses to landscape fragmentation: Implications for management strategies. Evolutionary Applications, 3, 291–304.

Bowcock, A. M., Ruizlinares, A., Tomfohrde, J., Minch, E., Kidd, J. R., & Cavallisforza, L. L. (1994). High-resolution of human evolutionary trees with polymorphic microsatellites. Nature, 368, 455–457.

Breiman, L. (2001). Random forests. Machine Learning, 45, 5–32.

Broquet, T., & Petit, E. J. (2009). Molecular Estimation of Dispersal for Ecology and Population Genetics. Annual Review of Ecology, Evolution, and Systematics, 40, 193–216.

Cavalli-Sforza, L. L., & Edwards, A. W. (1967). Phylogenetic analysis. Models and estimation procedures. American Journal of Human Genetics, 19, 233.

Cayuela, H., Rougemont, Q., Prunier, J. G., Moore, J.-S., Clobert, J., Besnard, A., & Bernatchez, L. (2018). Demographic and genetic approaches to study dispersal in wild animal populations: A methodological review. Molecular Ecology, 27, 3976–4010.

Coleman, R. A., Gauffre, B., Pavlova, A., Beheregaray, L. B., Kearns, J., Lyon, J., … Sunnucks, P. (2018). Artificial barriers prevent genetic recovery of small isolated populations of a low-mobility freshwater fish. Heredity. https://doi.org/10.1038/s41437-017-0008-3

Cooke, S. J., & Hinch, S. G. (2013). Improving the reliability of fishway attraction and passage efficiency estimates to inform fishway engineering, science, and practice. Ecological Engineering, 58, 123–132.

Couto, T. B., & Olden, J. D. (2018). Global proliferation of small hydropower plants—Science and policy. Frontiers in Ecology and the Environment. https://doi.org/10.1002/fee.1746

Csilléry, K., Blum, M. G., Gaggiotti, O. E., & François, O. (2010). Approximate Bayesian computation (ABC) in practice. Trends in Ecology & Evolution, 25, 410–418.

Davey, J. W., Hohenlohe, P. A., Etter, P. D., Boone, J. Q., Catchen, J. M., & Blaxter, M. L. (2011). Genome-wide genetic marker discovery and genotyping using next-generation sequencing. Nature Reviews Genetics, 12, 499–510.

Dudgeon, D., Arthington, A. H., Gessner, M. O., Kawabata, Z.-I., Knowler, D. J., Lévêque, C., … Sullivan, C. A. (2006). Freshwater biodiversity: Importance, threats, status and conservation challenges. Biological Reviews, 81, 163–182.

Fredrich, F., Ohmann, S., Curio, B., & Kirschbaum, F. (2003). Spawning migrations of the chub in the River Spree, Germany. Journal of Fish Biology, 63, 710–723.

Gauffre, B., Estoup, A., Bretagnolle, V., & Cosson, J. F. (2008). Spatial genetic structure of a small rodent in a heterogeneous landscape. Molecular Ecology, 17, 4619–4629.

Genuer, R., Poggi, J.-M., Tuleau-Malot, C., & Villa-Vialaneix, N. (2017). Random Forests for Big Data. Big Data Research, 9, 28–46.

Gibson, L., Wilman, E. N., & Laurance, W. F. (2017). How Green is =Green’ Energy? Trends in Ecology & Evolution, 32, 922–935.

Goudet, J. (1995). FSTAT (Version 1.2): A Computer Program to Calculate F-Statistics. Journal of Heredity, 86, 485–486.

Gouskov, A., Reyes, M., Wirthner-Bitterlin, L., & Vorburger, C. (2016). Fish population genetic structure shaped by hydroelectric power plants in the upper Rhine catchment. Evolutionary Applications, 9, 394–408.

Hague, M. T. J., & Routman, E. J. (2016). Does population size affect genetic diversity? A test with sympatric lizard species. Heredity, 116, 92–98.

Hand, B. K., Flint, C. G., Frissell, C. A., Muhlfeld, C. C., Devlin, S. P., Kennedy, B. P., … Stanford, J. A. (2018). A social-ecological perspective for riverscape management in the Columbia River Basin. Frontiers in Ecology and the Environment, 16, S23–S33.

Hawkins, P. R., Hortle, K. G., Phommanivong, S., & Singsua, Y. (2018). Underwater video monitoring of fish passage in the Mekong River at Sadam Channel, Khone Falls, Laos. River Research and Applications, 34, 232–243.

Hedrick, P. W. (2005). A Standardized Genetic Differentiation Measure. Evolution, 59, 1633–1638.

Jansson, R., Nilsson, C., & Malmqvist, B. (2007). Restoring freshwater ecosystems in riverine landscapes: The roles of connectivity and recovery processes. Freshwater Biology, 52, 589–596.

Januchowski-Hartley, S. R., Diebel, M., Doran, P. J., & McIntyre, P. B. (2014). Predicting road culvert passability for migratory fishes. Diversity and Distributions, 20, 1414–1424.

Jombart, T., Devillard, S., & Balloux, F. (2010). Discriminant analysis of principal components: A new method for the analysis of genetically structured populations. BMC Genetics, 11, 94.

Jost, L. (2008). *G* _ST_ and its relatives do not measure differentiation. Molecular Ecology, 17, 4015–4026.

Junge, C., Museth, J., Hindar, K., Kraabøl, M., & Vøllestad, L. A. (2014). Assessing the consequences of habitat fragmentation for two migratory salmonid fishes. Aquatic Conservation: Marine and Freshwater Ecosystems, 24, 297–311.

Kimura, M. (1983). The Neutral Theory of Molecular Evolution. Cambridge University Press.

Klütsch, C. F. C., Maduna, S. N., Polikarpova, N., Forfang, K., Aspholm, P. E., Nyman, T., … Hagen, S. B. (2019). Genetic changes caused by restocking and hydroelectric dams in demographically bottlenecked brown trout in a transnational subarctic riverine system. Ecology and Evolution, ece3.5191.

Landguth, E. L., Cushman, S. A., Schwartz, M. K., McKELVEY, K. S., Murphy, M., & Luikart, G. (2010). Quantifying the lag time to detect barriers in landscape genetics. Molecular Ecology, 19, 4179–4191.

Li, Y.-C., Korol, A. B., Fahima, T., Beiles, A., & Nevo, E. (2002). Microsatellites: Genomic distribution, putative functions and mutational mechanisms: a review. Molecular Ecology, 11, 2453–2465.

Liaw, A., & Wiener, M. (2002). Classification and Regression by randomForest. R News, 2, 18–22.

Lowe, W. H., & Allendorf, F. W. (2010). What can genetics tell us about population connectivity? Molecular Ecology, 19, 3038–3051.

McLaughlin, R. L., Smyth, E. R. B., Castro-Santos, T., Jones, M. L., Koops, M. A., Pratt, T. C., & Vélez-Espino, L.-A. (2013). Unintended consequences and trade-offs of fish passage. Fish and Fisheries, 14, 580–604.

McNeish, D. M. (2014). Analyzing Clustered Data with OLS Regression: The Effect of a Hierarchical Data Structure. 40, 6.

McRae, B. H. (2006). Isolation by resistance. Evolution, 60, 1551–1561.

Meirmans, P. G. (2006). Using the Amova Framework to Estimate a Standardized Genetic Differentiation Measure. Evolution, 60, 2399–2402.

Meirmans, P. G., & Hedrick, P. W. (2011). Assessing population structure: FST and related measures. Molecular Ecology Resources, 11, 5–18.

Neuenschwander, S., Michaud, F., & Goudet, J. (2019). QuantiNemo 2: A Swiss knife to simulate complex demographic and genetic scenarios, forward and backward in time. Bioinformatics, 35, 886–888.

Nilsson, C. (2005). Fragmentation and Flow Regulation of the World’s Large River Systems. Science, 308, 405–408.

Paz-Vinas, I., Comte, L., Chevalier, M., Dubut, V., Veyssiere, C., Grenouillet, G., … Blanchet, S. (2013). Combining genetic and demographic data for prioritizing conservation actions: Insights from a threatened fish species. Ecology and Evolution, 3, 2696–2710.

Poff, N. L., & Schmidt, J. C. (2016). How dams can go with the flow. Science (New York, N.Y.), 353, 1099–1100.

Poulet, N. (2007). Impact of weirs on fish communities in a piedmont stream. River Research and Applications, 23, 1038–1047.

Pracheil, B. M., Mestl, G. E., & Pegg, M. A. (2015). Movement through Dams Facilitates Population Connectivity in a Large River. River Research and Applications, 31, 517–525.

Pritchard, J. K., Stephens, M., & Donnelly, P. (2000). Inference of population structure using multilocus genotype data. Genetics, 155, 945–959.

Prunier, J. G., Dubut, V., Chikhi, L., & Blanchet, S. (2017). Contribution of spatial heterogeneity in effective population sizes to the variance in pairwise measures of genetic differentiation. Methods in Ecology and Evolution, 8, 1866–1877.

Prunier, J. G., Dubut, V., Loot, G., Tudesque, L., & Blanchet, S. (2018). The relative contribution of river network structure and anthropogenic stressors to spatial patterns of genetic diversity in two freshwater fishes: A multiple-stressors approach. Freshwater Biology, 63, 6–21.

R Development Core Team. (2014). R: A Language and Environment for Statistical Computing, R Foundation for Statistical Computing. Retrieved from http://www.R-project.org

Raeymaekers, J. A. M., Raeymaekers, D., Koizumi, I., Geldof, S., & Volckaert, F. A. M. (2009). Guidelines for restoring connectivity around water mills: A population genetic approach to the management of riverine fish. Journal of Applied Ecology, 46, 562–571.

Reid, A. J., Carlson, A. K., Creed, I. F., Eliason, E. J., Gell, P. A., Johnson, P. T. J., … Cooke, S. J. (2018). Emerging threats and persistent conservation challenges for freshwater biodiversity. Biological Reviews. https://doi.org/10.1111/brv.12480

Richardson, J. L., Brady, S. P., Wang, I. J., & Spear, S. F. (2016). Navigating the pitfalls and promise of landscape genetics. Molecular Ecology, 25, 849–863.

Rousset, F. (2008). GENEPOP ’007: A complete re-implementation of the GENEPOP software for Windows and Linux. Molecular Ecology Resources, 8, 103–106.

Roy, S. G., Uchida, E., de Souza, S. P., Blachly, B., Fox, E., Gardner, K., … Hart, D. (2018). A multiscale approach to balance trade-offs among dam infrastructure, river restoration, and cost. Proceedings of the National Academy of Sciences, 115, 12069–12074.

Saint-Pé, K., Blanchet, S., Tissot, L., Poulet, N., Plasseraud, O., Loot, G., … Prunier, J. G. (2018). Genetic admixture between captive-bred and wild individuals affects patterns of dispersal in a brown trout (*Salmo trutta*) population. Conservation Genetics, 19, 1269–1279.

Schlötterer, C. (2000). Evolutionary dynamics of microsatellite DNA. Chromosoma, 109, 365–371.

Selkoe, K. A., Scribner, K. T., & Galindo, H. M. (2015). Waterscape Genetics—Applications of Landscape Genetics to Rivers, Lakes, and Seas. In N. Balkenhol, S. A. Cushman, A. T. Storfer, & L. P. Waits (Eds.), Landscape Genetics (pp. 220–246). Chichester, UK: John Wiley & Sons, Ltd.

Silva, A. T., Lucas, M. C., Castro-Santos, T., Katopodis, C., Baumgartner, L. J., Thiem, J. D., … Cooke, S. J. (2018). The future of fish passage science, engineering, and practice. Fish and Fisheries, 19, 340–362.

Song, C., Omalley, A., Roy, S. G., Barber, B. L., Zydlewski, J., & Mo, W. (2019). Managing dams for energy and fish tradeoffs: What does a win-win solution take? Science of The Total Environment, 669, 833–843.

Städele, V., & Vigilant, L. (2016). Strategies for determining kinship in wild populations using genetic data. Ecology and Evolution, 6, 6107–6120.

Storfer, A., Murphy, M. A., Spear, S. F., Holderegger, R., & Waits, L. P. (2010). Landscape genetics: Where are we now? Molecular Ecology, 19, 3496–3514.

Sundqvist, L., Keenan, K., Zackrisson, M., Prodöhl, P., & Kleinhans, D. (2016). Directional genetic differentiation and relative migration. Ecology and Evolution, n/a–n/a.

Turgeon, K., Turpin, C., & Gregory-Eaves, I. (2019). Dams have varying impacts on fish communities across latitudes: A quantitative synthesis. Ecology Letters, ele.13283.

Wang, J. L. (2005). Estimation of effective population sizes from data on genetic markers. Philosophical Transactions of the Royal Society B-Biological Sciences, 360, 1395–1409.

Weir, B. S., & Cockerham, C. C. (1984). Estimating F-Statistics for the analysis of population structure. Evolution, 38, 1358–1370.

Wilson, G. A., & Rannala, B. (2003). Bayesian inference of recent migration rates using multilocus genotypes. Genetics, 163, 1177–1191.

Winter, D. J. (2012). MMOD: An R library for the calculation of population differentiation statistics. Molecular Ecology Resources, 12, 1158–1160.

Yue, G. H., David, L., & Orban, L. (2007). Mutation rate and pattern of microsatellites in common carp (Cyprinus carpio L.). Genetica, 129, 329–331.

