## Appendices for "Quantifying the individual impact of artificial barriers in freshwaters: A standardized and absolute genetic index of fragmentation"

Running title: Standardized genetic index of fragmentation

Jérôme G. Prunier [1] *, Camille Poesy [1], Vincent Dubut [2], Charlotte Veyssière [3], Géraldine Loot [3], Nicolas Poulet [4] & Simon Blanchet [1,3]*

[1] Centre National de la Recherche Scientifique (CNRS), Université Paul Sabatier (UPS); Station d'Ecologie Théorique et Expérimentale, UMR 5321, F-09200 Moulis, France

[2] Aix Marseille Univ, CNRS, IRD, Avignon Univ, IMBE, Marseille, France

[3] CNRS, UPS, École Nationale de Formation Agronomique (ENFA) ; UMR 5174 EDB (Laboratoire Évolution & Diversité Biologique), 118 route de Narbonne, F-31062 Toulouse cedex 4, France

[4] Pôle écohydraulique AFB-IMT, allée du Pr Camille Soula, 31400 Toulouse, France

*Corresponding authors:

Jérôme G. Prunier and Simon Blanchet

Station d'Ecologie Théorique et Expérimentale, UMR 5321, F-09200 Moulis, France

2 route du CNRS, 09200 Moulis

Phone: (+33)561040361

### Appendix S1: The FINDEX() R-function: Walkthrough for the computation of the $F_{INDEX}$

The $F_{INDEX}$ computation was automated within a user-friendly R-function. The FINDEX() R-function is embedded within the "FINDEXpackage.rda" file (http://www.jeromeprunier.fr/Tools.html). This .rda file also contains all RandomForest predictions required to compute the $F_{INDEX}$. Users are invited to download this .rda file within their working directory, and to install required R-libraries ("adegenet", "randomForest", "mmod", "lme4" and "reshape2").

```
# Installing required libraries (if needed)
    install.packages(c("adegenet ", " randomForest ","mmod","lme4","reshape2"),dependencies=TRUE)

# Going to working directory (where " FINDEXpackage.rda" is located)
    setwd('path_to_working_directory')

# Load .rda file to import the FINDEX() R-function and all other required files in the R-environment
    load("FINDEXpackage.rda")
```

Users are then simply expected to provide one or several empirical genotypic datasets (in the *genepop* format with a ".gen" extension) and a parameter file (a dataframe in ".txt" format) with a line for each combination of both an obstacle and a species and at least **eight mandatory columns**:

- "**Species**": the name of the considered species (factor);
- "**Barrier**": the name of the considered obstacle (factor);
- "**Upstream**": the name of the upstream population as found in the corresponding *genepop* file (factor);
- "**Downstream**": the name of the downstream population as found in the corresponding *genepop* file (factor);
- "**Creation**": the date of barrier creation (numeric);
- "**Generations**": the number of generations elapsed since barrier creation (numeric; this number of generations is to be estimated from the life-history traits of the considered species);
- "**Genepop**": the name of the *genepop* file from which genotypes are to be extracted (with or without the ".gen" extension); see https://genepop.curtin.edu.au/help_input.html#Input for help with the formatting of *genepop* files;
- "**Digits**": the number of characters used to code an allele in the *genepop* file.

Note that the parameter file may contain any additional column, provided that these columns add information about **barriers** (for instance, height, spatial localization, etc.).

In an illustrative purpose, we provide a parameter file "*Prunier2018_illustration.txt*" as well as two *genepop* files ("*Phoxinus.gen*" and "*Gobio.gen*") to test the FINDEX R-function. These files are provided as a compressed zip-file (http://www.jeromeprunier.fr/Tools.html).
The "*Prunier2018_illustration.txt*" looks like the following:

| River | **Species** | **Barrier** | **Upstream** | **Downstream** | **Creation** | **Generations** | **Genepop** | **Digits** |
|---|---|---|---|---|---|---|---|---|
| Cele | Gobio | SCC | SCCup | SCCdown | 1960 | 20 | Gobio.gen | 3 |
| Viaur | Gobio | CAP | CAPup | CAPdown | 1700 | 124 | Gobio | 3 |
| Cele | Phoxinus | SCC | SCCup29 | SCCdown30 | 1960 | 25 | Phoxinus.gen | 3 |
| Viaur | Phoxinus | CAP | CAPup30 | CAPdown30 | 1700 | 155 | Phoxinus | 3 |

It contains the eight mandatory columns (in bold) and an additional column indicating the river in which each barrier is located.
In the parameter file, the names of *genepop* files (Genepop column) can be provided with or without the .gen extension.

In the “*Phoxinus.gen*” file, genotypes are provided with the **individual numbering** directly attached to the population name (e.g., SCCup**01**, CAPdown**02**, etc.). In that case, the name of each population (Upstream and Downstream columns) is the name of the last genotype in that population (e.g., SCCup29, CAPdown30, etc.).
In the “*Gobio.gen*” file, genotypes are either provided with the **individual numbering** separated from the population name by an underscore (e.g., SCCup**_01**, SCCdown**_02**, etc.), in which case the name of each population (Upstream and Downstream columns) is the part of the genotype name located before the underscore (e.g., SCCup, SCCdown), or without any individual numbering, in which case the name of each population (Upstream and Downstream columns) is directly the name of genotypes.

Once these illustration files have been extracted within the working directory, users simply have to run the FINDEX() R-function as follows:

```
# Running the FINDEX() function on illustration files
    FINDEX_test=FINDEX(input_file="Prunier2018_illustration.txt”)
```

A “verbose” option can be turned off (“verbose=FALSE”), in which case no information about the computation progress will be provided.
This command will launch the computation of the $F_{INDEX}$ and return a list with two elements: “output1” and “output2”.
“output1” is a dataframe providing results for each combination of both an obstacle and a species, with at least the following columns (as well as any additional (non-mandatory) column from the input file):

- “Species”: the name of the considered species (factor)
- “Barrier”: the name of the considered obstacle (factor)
- “Upstream”: the name of the upstream population (factor)
- “Dowstream”: the name of the downstream population (factor)
- “Creation”: the date of barrier creation (numeric).
- “Generations”: the number of generations elapsed since barrier creation (numeric)
- “He_Up”: the expected heterozygosity in the upstream population (numeric)
- “He_Down”: the expected heterozygosity in the downstream population (numeric)
- “He”: the averaged expected heterozygosity across populations (numeric)
- “Obs_PhiST”: the observed measure of ϕ’st (numeric)
- “Obs_GST2”: the observed measure of G’’st (numeric)
- “Findex”: the computed $F_{INDEX}$ (numeric)
- “ciFindex” : the 95% confidence interval about the $F_{INDEX}$ (numeric)

By default, “output2” is a dataframe providing results **for each obstacle** with $F_{INDEX}$ values averaged across species, with at least the following columns:

- “Barrier”: the name of the considered obstacle (factor)
- “Creation”: the date of barrier creation (numeric)
- “Findex”: the computed $F_{INDEX}$ (numeric)
- “ciFindex” : the 95% confidence interval about $F_{INDEX}$ (numeric)

The FINDEX() function can also be used to compile $F_{INDEX}$ values across other factors than obstacles (the default). In that case, users will have to use the “CompileBy” option and indicate which additional column from the input parameter file will be used for averaging. This column must be coded as factors. From the illustration files, $F_{INDEX}$ values could for instance be compiled across rivers, as follows:

```
# Running the FINDEX() function on illustration files with Findex values compiled across rivers
    FINDEX_test=FINDEX(input_file="Prunier2018_illustration.txt”, CompileBy=”River”)
```

Here, the "output2" dataframe will provide results for each river, with $F_{INDEX}$ values averaged across both obstacles and species.

Eventually, the results can be plotted using the following commands:

```
# Loading required libraries (to be installed if needed)
    require("ggplot2")
    require("viridis")

# Plotting results for each combination of both an obstacle and a species
    ggplot(FINDEX_test$output1, aes(x=reorder(Barrier, as.numeric(FINDEX_test$output1$Creation)), y=
            Findex,group=Species, fill= Findex))+
        geom_bar(width = 0.9,stat="identity", position=position_dodge(.9))+
        geom_text(aes(y = 105, label = Species, angle=0),position = position_dodge(width = .9))+
        coord_cartesian(ylim=c(0,105)) +
        geom_errorbar(aes(ymin= Findex-ciFindex, ymax= Findex+ciFindex), width=0,
            position=position_dodge(.9))+
        scale_fill_viridis(discrete = FALSE,begin = 0.1,end=0.9,limits=c(0,100))+
        geom_hline(yintercept=c(0,20,90,100), linetype="dashed", color = "black")+
        scale_x_discrete(name ="Obstacles")


# Plotting results for each obstacle with FINDEX averaged across species
    ggplot(FINDEX_test$output2, aes(x=reorder(Barrier, as.numeric(FINDEX_test$output2$Creation)), y=
            Findex,group=Barrier, fill= Findex)) +
        geom_bar(width = 0.9,stat="identity", position=position_dodge(preserve = 'single'))+
        coord_cartesian(ylim=c(0,105)) +
        geom_errorbar(aes(ymin= Findex-ciFindex, ymax= Findex+ciFindex), width=0.1,
            position=position_dodge(width = 0,preserve = 'single'))+
        scale_fill_viridis(discrete = FALSE,begin = 0.1,end=0.9,limits=c(0,100))+
        geom_hline(yintercept=c(0,20,90,100), linetype="dashed", color = "black")+
        scale_x_discrete(name ="Obstacles")
```

Note that users also have the possibility to provide their own $GD_{min}$ and $GD_{max}$ values, in the form of two dataframes, each with two columns ('GST2' and 'PhiST') and the same number of rows as the input file (NA values are allowed, in which case the predicted $GD_{min}$ and/or $GD_{max}$ values would be used). These options are yet to be used with caution (see main text for details).

```
# Creating GDmin and GDmax dataframes and running the FINDEX() function
  GDMIN=data.frame(GST2= rep(0.01,4),PhiST= rep(0.02,4))
  GDMAX=data.frame(GST2=rep(0.2,4),PhiST= rep(0.3,4))
  FINDEX_test=FINDEX(input_file="Prunier2018_illustration.txt",GDmin=GDMIN,GDmax=GDMAX)
```

### Appendix S2

When considering two populations separated by an obstacle whose impact on gene flow is unknown, simulations can be used to predict the expected range of any measure $k$ of genetic differentiation $GD^k$ given the age of the obstacle (expressed in number of generations since barrier creation), the averaged expected heterozygosity of the two populations (a proxy for effective population size; see Appendix S5; Hague & Routman, 2016; Prunier et al., 2017) and the simulated impact of the obstacle on gene flow. When the obstacle has no impact on gene flow, then $GD^k = GD^k_{min}$ , with $GD^k_{min}$ the non-null level of genetic differentiation that may be expected under the sole effect of genetic drift, mutations and random sampling. When the obstacle is a total barrier to gene flow, then $GD^k = GD^k_{max}$. The actual observed measure of genetic differentiation $GD^k_{obs}$ can then be set to $GD^k_{min}$ when $GD^k_{obs} < GD^k_{min}$ or to $GD^k_{max}$ when $GD^k_{obs} > GD^k_{max}$

Mathematically, and considering any pairwise measure of genetic differentiation as a *relative risk* ("risk" for two populations to be genetically distinct), dividing $GD_{obs}$ by $GD_{max}$ amounts to computing a *risk-ratio* (Borenstein, 2009): a risk-ratio of 50% would for instance mean that the *risk* of being genetically distinct in the observed situation is 0.5 times the *risk* of being genetically distinct in the worst-case scenario (total genetic differentiation); similarly, a risk-ratio of 100% would mean that the *risk* of being genetically distinct in the observed situation is as high as the *risk* of being genetically distinct in the worst-case scenario (total genetic differentiation). For risk ratios, computations are usually carried out on a log-scale. A standardized estimate $c^k_{obs}$ of the amount of gene flow that gets through the obstacle when compared to the worst-case scenario can thus be computed as the inverse of the log-risk-ratio between $GD^k_{obs}$ and $GD^k_{max}$ :

$$c^k_{obs} = -\,ln\left(\frac{GD^k_{obs}}{GD^k_{max}}\right)$$

With $GD^k_{obs}$ ranging from $GD^k_{min}$ to $GD^k_{max}$, this index ranges from $c^k_{min} = -ln\left(\frac{GD^k_{max}}{GD^k_{max}}\right) = -\ln(1) = 0$ to $c^k_{max} = -ln\left(\frac{GD^k_{min}}{GD^k_{max}}\right) > 0$. It can thus be linearly rescaled to range from 0 to 1 by dividing it by $c^k_{max}$, leading to:

$$C^k_{index} = \frac{c^k_{obs}}{c^k_{max}} = \frac{-ln\left(\frac{GD^k_{obs}}{GD^k_{max}}\right)}{-ln\left(\frac{GD^k_{min}}{GD^k_{max}}\right)} = \frac{ln\left(GD^k_{obs}\right) - ln(GD^k_{max})}{ln(GD^k_{min}) - ln(GD^k_{max})}$$

The $C^k_{index}$ is an index of connectivity: it is simply the ratio of the observed log-risk-ratio (numerator) to the log-risk-ratio expected in the absence of any barrier to gene flow (denominator). The proposed index of fragmentation $F$ (eqn. 1 in main text) is finally computed as follows:

$$F^k_{index} = 1 - C^k_{index} = 1 - \frac{ln\left(GD^k_{obs}\right) - ln(GD^k_{max})}{ln(GD^k_{min}) - ln(GD^k_{max})} = \frac{ln\left(GD^k_{min}\right) - ln\left(GD^k_{obs}\right)}{ln(GD^k_{min}) - ln(GD^k_{max})}$$

### Appendix S3

To investigate the sensitivity of various metrics of genetic differentiation to sample size, we used the program QuantiNemo2 (Neuenschwander et al., 2019) to simulate gene flow between two adjacent demes of size 2000 over 400 non-overlapping generations. Each deme was initiated with 2000 individuals and kept at a constant size over generations. Genetic polymorphism was based on 15 microsatellite loci and 20 alleles per locus. The mutation rate $\mu$, following a stepwise mutation model, was set to $5\times10^{-4}$. Genotypes were randomly assigned at the beginning of simulations and the crossing rate was set to 0.5. We ran ten simulation replicates and then considered 98 sample sizes (from 30 to 1000 individuals sampled without replacement in each population), resulting in a total of 980 simulated genetic datasets in the *Fstat* format (Goudet, 1995), further converted into the *genepop* format (Rousset, 2008) using R (R Development Core Team, 2014). For each dataset, we computed the distance based on the proportion of shared alleles (Bowcock et al., 1994) using the R-package *PopGenReport* (Adamack & Gruber, 2014), the Weir and Cockerham's θst (Weir & Cockerham, 1984), the Cavalli-Sforza and Edwards' Chord distance (Cavalli-Sforza & Edwards, 1967), the Nei's minimum distance (Nei, 1973b), the Rogers' distance (Rogers, 1972), the Sanghvi's distance (Sanghvi, 1953) and the Prevosti et al. distance (Prevosti et al., 1975) using the R-package *hierfstat* (Goudet, 2005); the Nei's Gst (Nei, 1973a; Nei & Chesser, 1983), the Hedrick's G''st (Hedrick, 2005; Meirmans & Hedrick, 2011), the Meirmans' ϕ'st (Meirmans, 2006) and the Jost's D (2008) using the R-package *mmod* (Winter, 2012).

We then plotted mean genetic distances (± standard deviation) computed over the 10 replicates against sample size. Several measures of genetic distances were found to be sensitive to sample size, as shown in figure S3a. All the following metrics indeed showed a systematic increase when sample size decreased: the distance based on the proportion of shared alleles, the Cavalli-Sforza and Edwards' Chord distance, the Nei's minimum distance, the Rogers' distance, the Sanghvi's distance and the Prevosti et al. distance. Furthermore, all these metrics but the Cavalli-Sforza and Edwards' Chord distance and the Nei's minimum distance showed non-null values whatever the sample size, despite total gene flow. On the contrary, and as shown in Figure S3b, the following metrics did not show any systematic increase with the decrease in sample size and were thus retained for further investigations (see Appendix S4): the Weir and Cockerham's θst, the Nei's Gst, the Hedrick's G''st, the Meirmans' ϕ'st and the Jost's D (2008).

**Figure S3:** Mean genetic distances (± standard deviation) computed over the 10 replicates and plotted against sample size. In panel A, six metrics show a systematic bias towards 1 as sample size decreases, contrary to the five retained metrics shown in panel B.

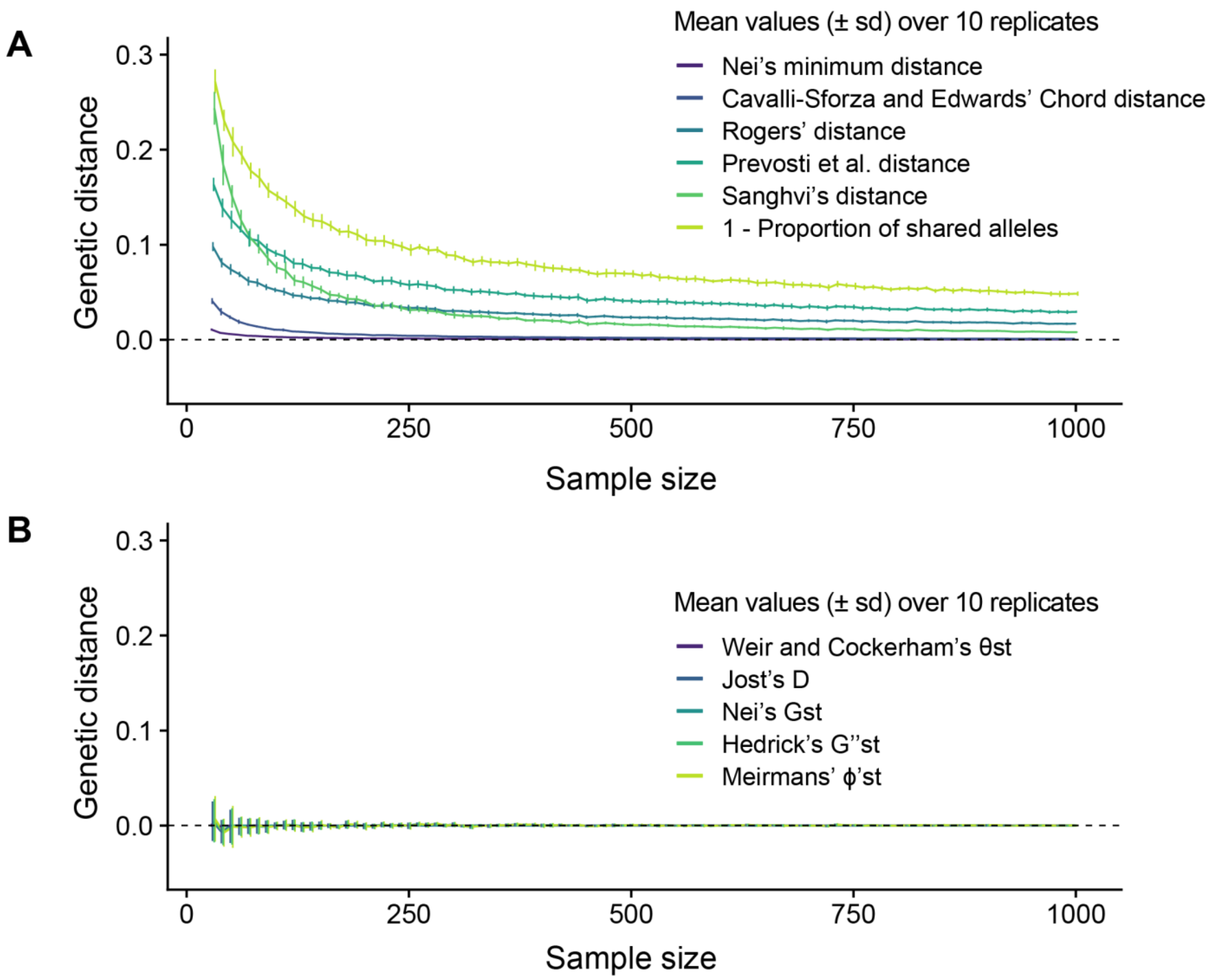

### Appendix S4

To investigate the behavior of the five retained metrics of genetic differentiation (the Weir and Cockerham's θst, the Nei's Gst, the Hedrick's G''st, the Meirmans' ϕ'st and the Jost's D (2008); see Appendix S3), we used a subset of the genetic data simulated for $GD_{max}$ predictions (see main text for details). For each mutation rate $\mu$ ($5\times10^{-5}$ or $5\times10^{-4}$) and for effective population sizes of 50 or 2000, we plotted the mean genetic distances (± standard deviation) computed over the 20 replicates against the number of generations elapsed since the beginning of simulations. Barrier creation (crossing rate dropping from 0.5 to 0) occurred at generation 400 (Figure S4).

**Figure S4:** Mean value GD (± standard deviation) of five metrics of genetic differentiation computed over 20 replicates and plotted against the number of generations elapsed since the beginning of simulations (Gen) according to the effective population size (*Ne* = 30, left panels A and C; or *Ne* = 2000; right panels B and D) and the mutation rate $\mu$ ($5\times10^{-5}$, panels A and B; $5\times10^{-4}$, panels C and D). Barrier creation occurred at generation 400.

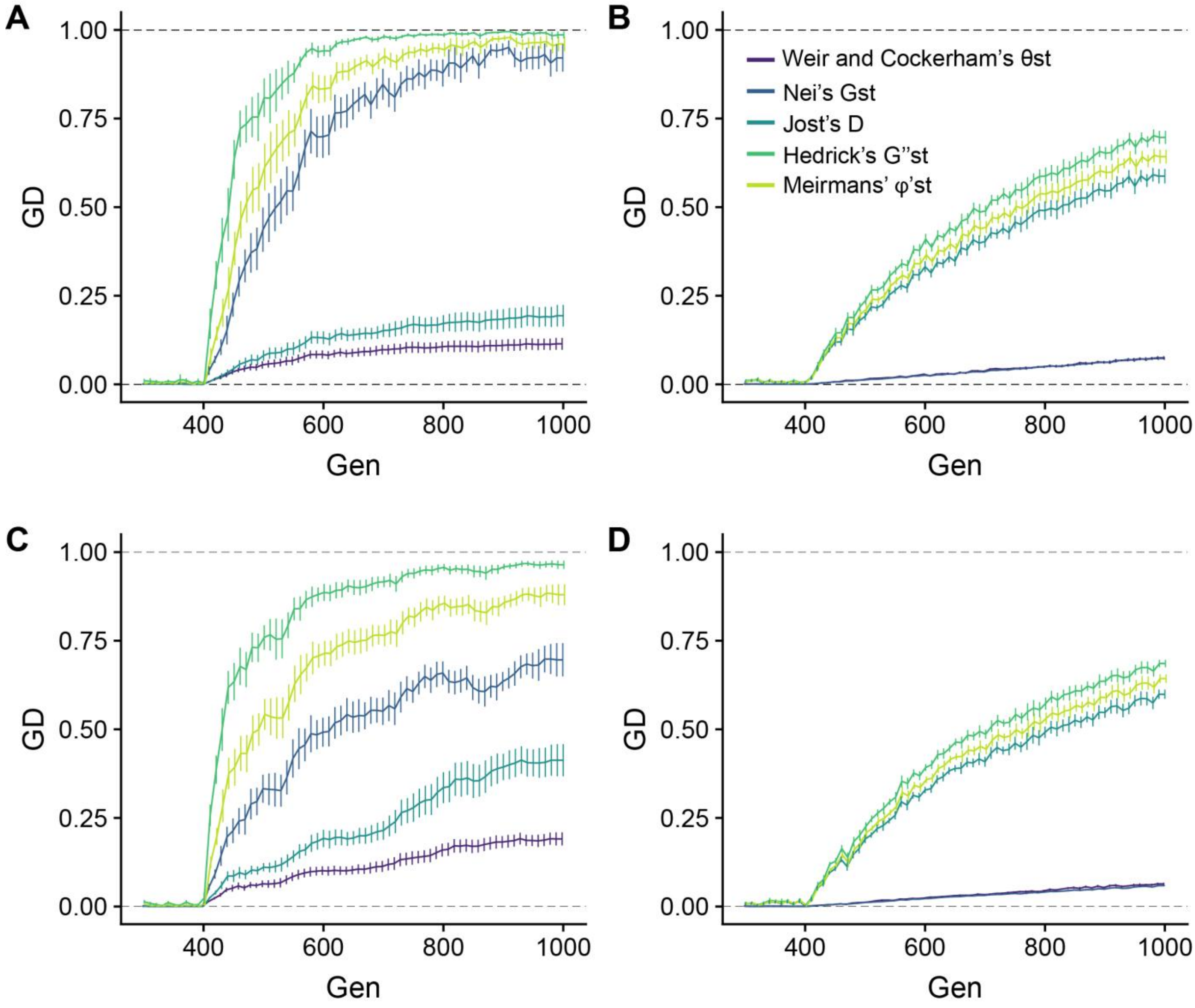

The five retained measures of genetic differentiation were null before barrier creation and quickly increased after barrier creation, whatever the effective population size (*Ne* = 30, left panels A and C; or *Ne* = 2000; right panels B and D) or the mutation rate $\mu$ ($5\times10^{-5}$, panels A and B; $5\times10^{-4}$, panels C and D). Nevertheless, two metrics were found to be highly sensitive to mutation rate when populations are of small effective sizes (Nei's Gst and Jost's D) and two metrics were found to show very little variability when populations are of large effective sizes (Nei's Gst and Weir and Cockerham's θst). We thus only retained the Hedrick's G''st and the Meirmans' ϕ'st as robust metrics of genetic differentiation.

### Appendix S5

When averaged over the two sampled populations (30 genotypes each) just before the creation of the barrier (at generation 400), expected heterozygosity *He* (computed using the R-package *adegenet*; Jombart, 2008) showed a positive monotonic relationship with carrying capacity, that is, with simulated effective population sizes *Ne* (see Figure S5a, as well as main text for details about simulations). Furthermore, *He* decreased after barrier creation, mimicking the expected decrease in *Ne* as the initial population is split into two adjacent yet disconnected sub-populations (see Figure S5b). As a consequence, *He* may be considered a good proxy for effective population sizes (Hague & Routman, 2016; Prunier et al., 2017).

**Figure S5a:** Mean expected heterozygosity against mean effective population size at generation 400, according to mutation rate.

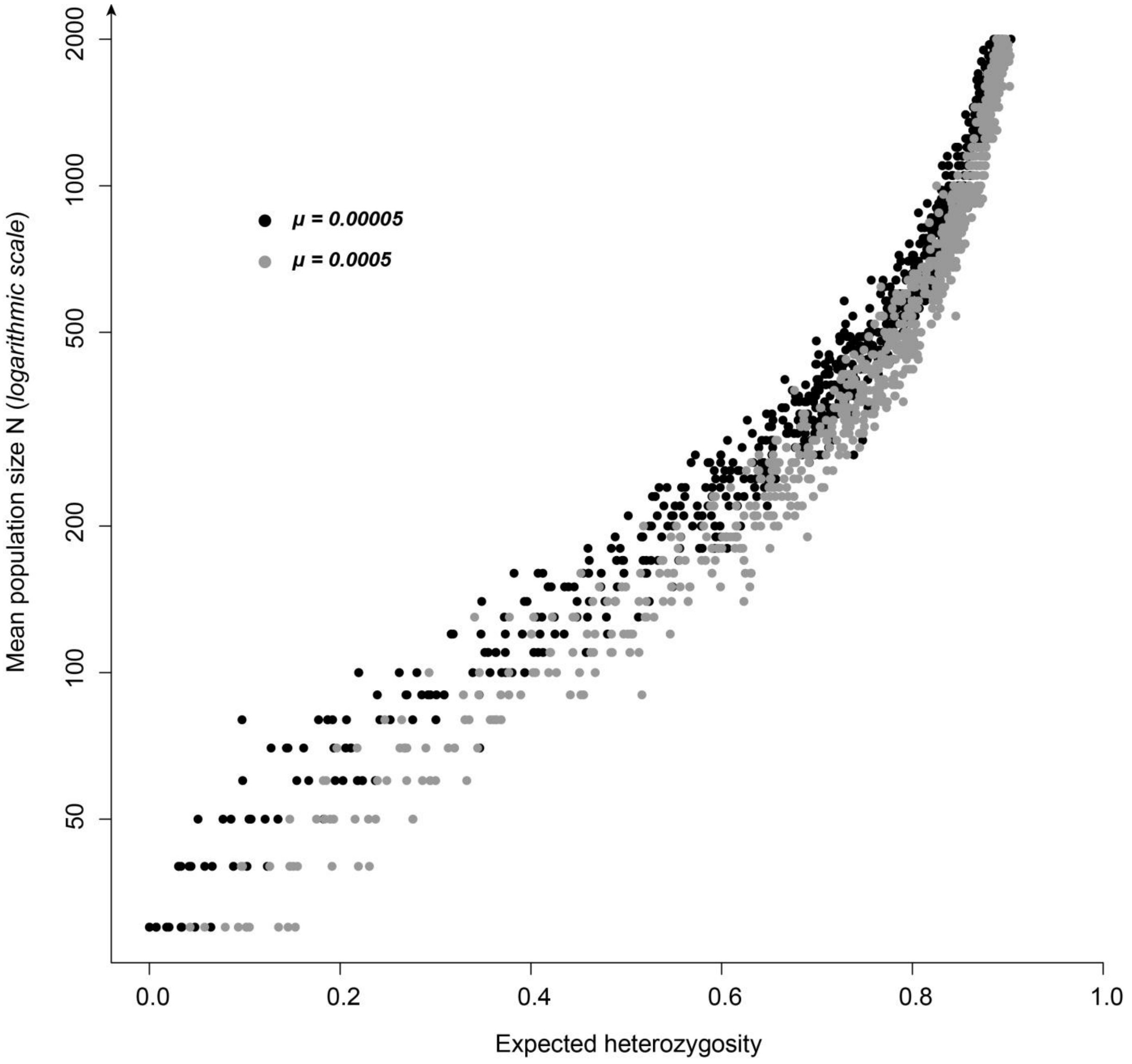

**Figure S5b:** For four levels of carrying capacity (K = 30, 100, 500 and 1000) and two mutation rates (μ = 0.0005 and 0.00005), evolution of mean expected heterozygosity over time since barrier creation. Note that the use of two mutation rates allowed simulating a large range of *He* values for each level of carrying capacity (with lower *He* values at a low mutation rate and higher *He* values at a high mutation rate).

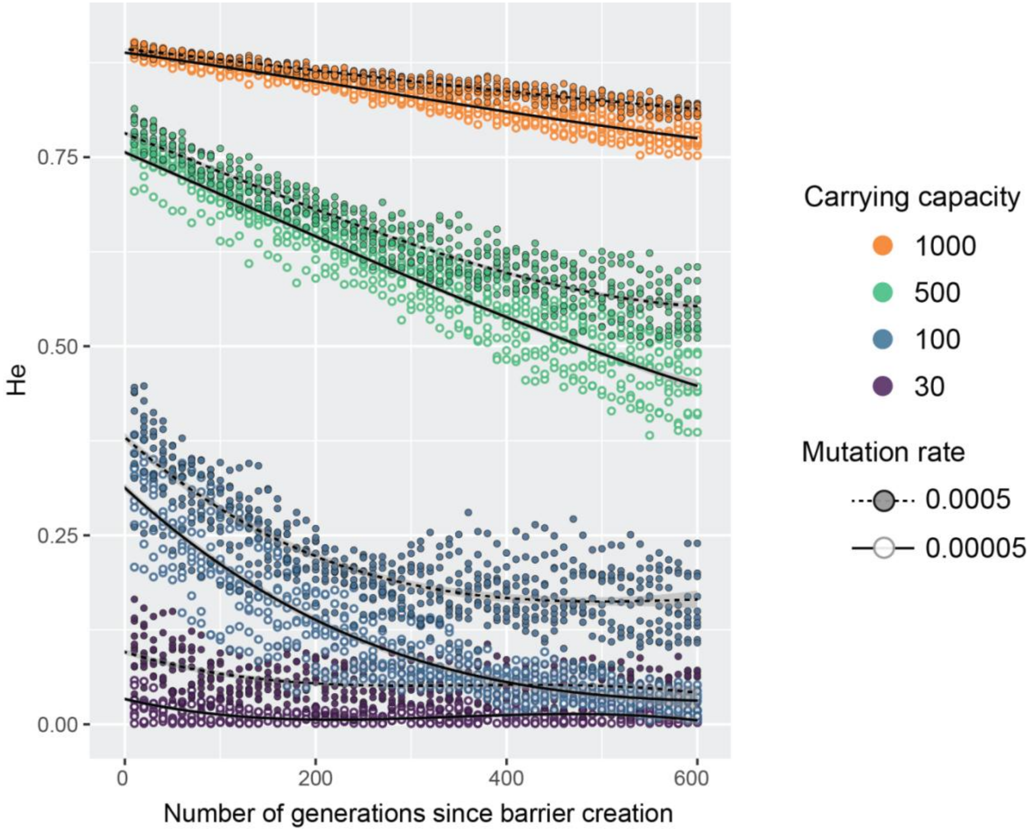

### Appendix S6

For each mutation rate $\mu$ ($\mu$ = 5.10$^{-5}$ or $\mu$ = 5.10$^{-4}$), distribution of measures of genetic differentiation GD (either G''st or φ'st) computed before the creation of the barrier. $GD_{min}$ values, indicated by vertical dashed lines, were computed as the fifth percentile of non-null simulated GD values.

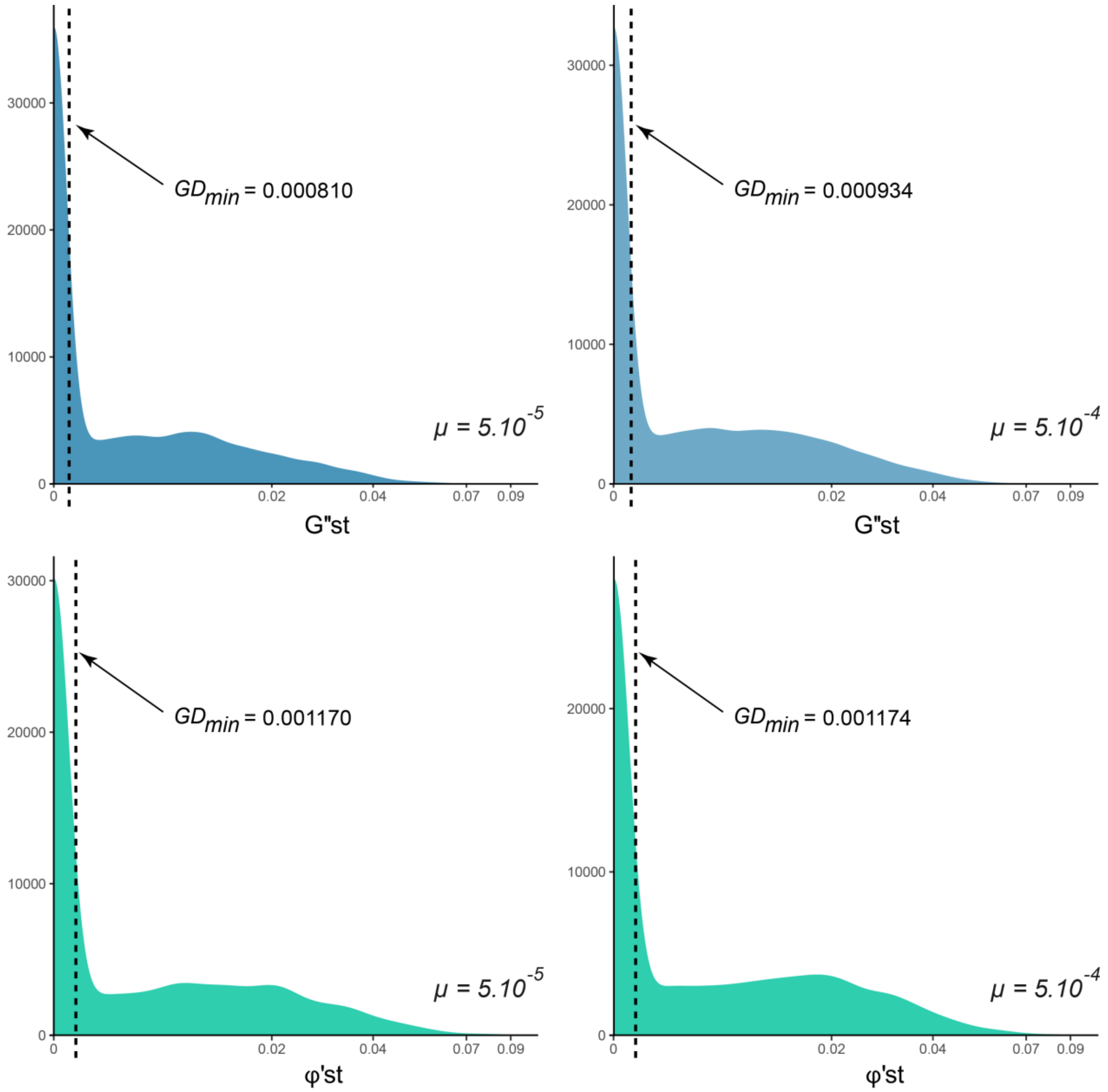

### Appendix S7

To validate the proposed genetic index of fragmentation $F_{INDEX}$ using realistic genetic datasets, each microsatellite locus was given a unique stepwise mutation rate $\mu$ randomly picked from a log-normal distribution with a mean of $5\times10^{-4}$ and a standard deviation of 2, thus ranging from $5\times10^{-5}$ to $5\times10^{-3}$ . The following figure illustrates the corresponding simulated distribution based on 10000 values. Red bars at the bottom of the histogram stand for the 15 randomly picked stepwise mutation rate $\mu$ used in Quantinemo simulations. These mutation rates ranged from $1.014\times10^{-5}$ to $2.181\times10^{-3}$.

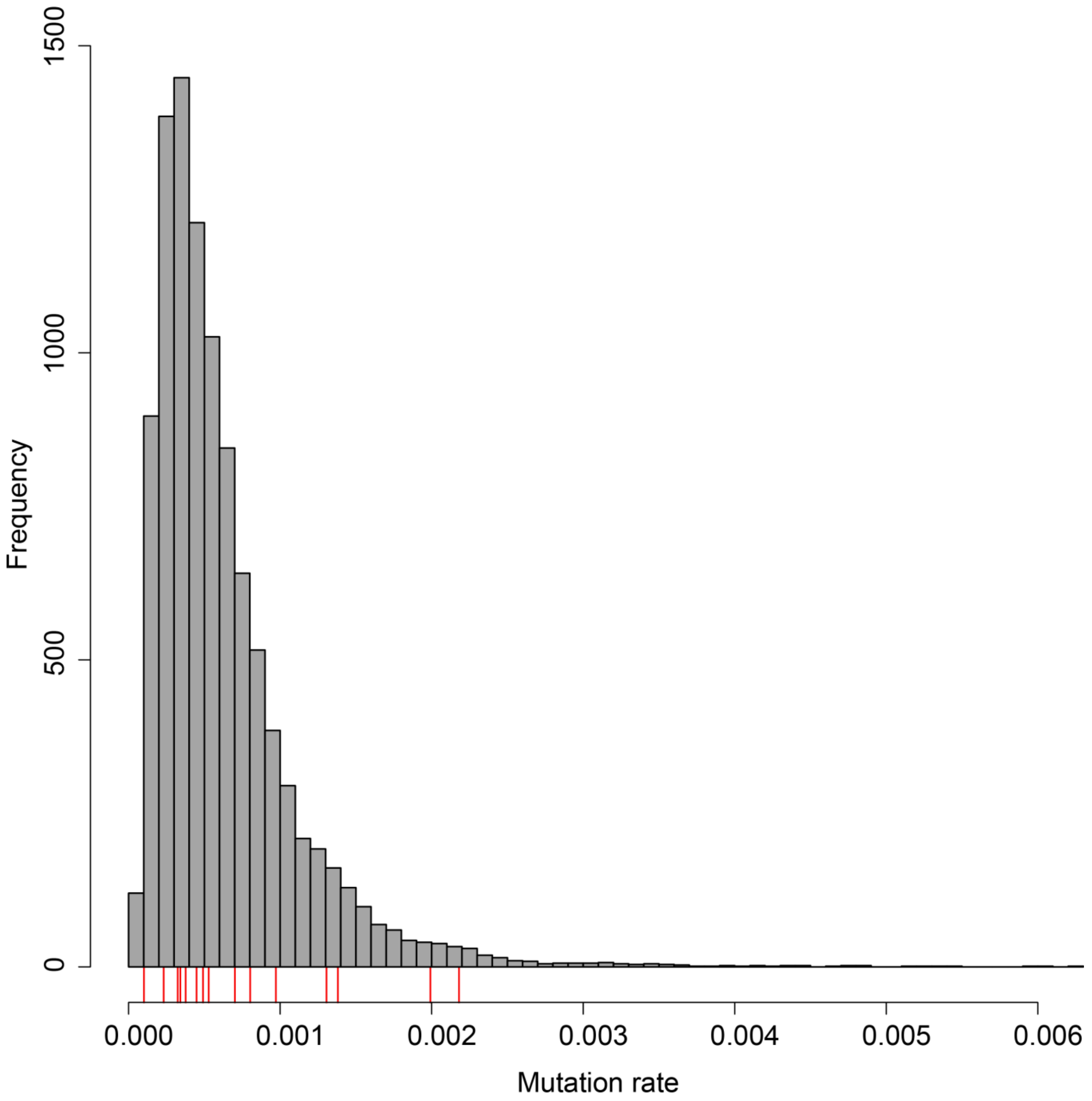

### Appendix S8

**Figure S8a:** Figure adapted from Gouskov et al. (2016). Localization of the six selected dams from the upper Rhine catchment. Bars represent barriers and the numbers correspond to the barrier numbers in Table 1 (main text). In red, dams identified as responsible for a decrease in gene flow according to the proposed genetic index of fragmentation $F_{INDEX}$.

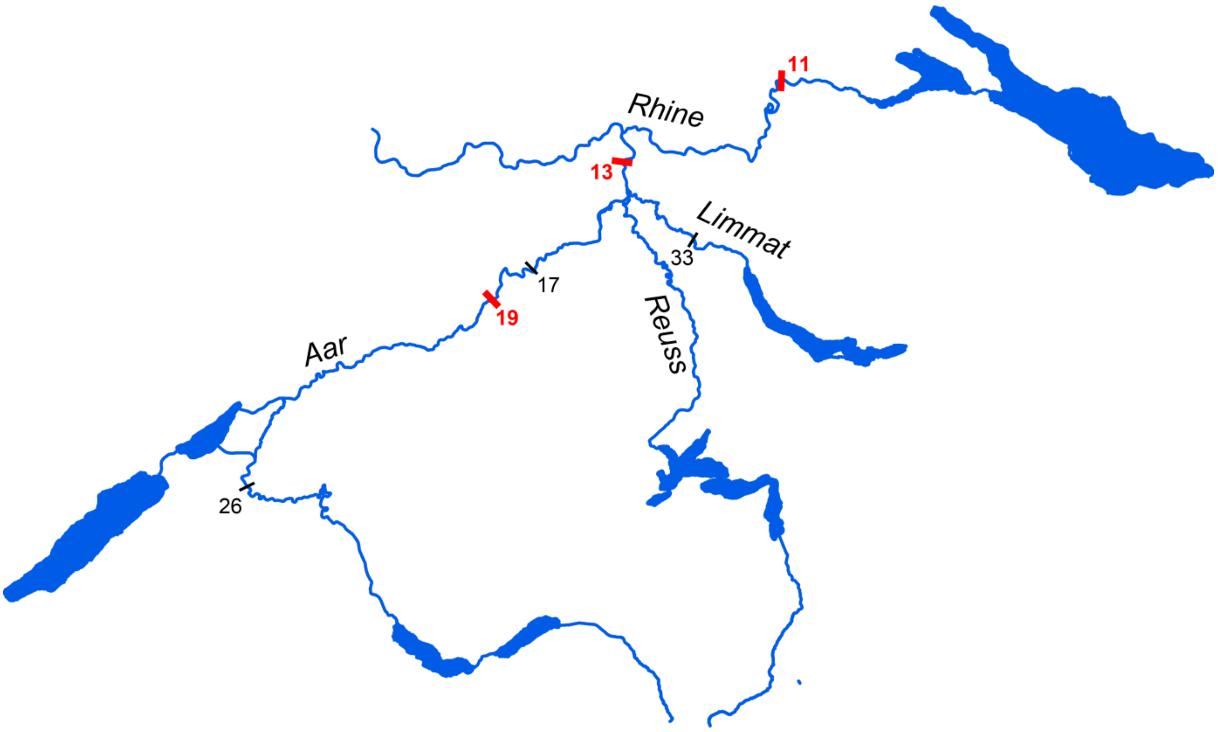

**Figure S8b:** Figure adapted from Prunier et al. (2018). Localization of the 8 selected weirs from the Célé and the Viaur Rivers. Triangles represent barriers, named as in Table 1 (main text). In blue, the only weir identified as responsible for a decrease in gene flow in both *Gobio occitaniae* and *Phoxinus phoxinus* according to the proposed genetic index of fragmentation $F_{INDEX}$. In red, weirs identified as responsible for a decrease in gene flow in *Gobio occitaniae* only.

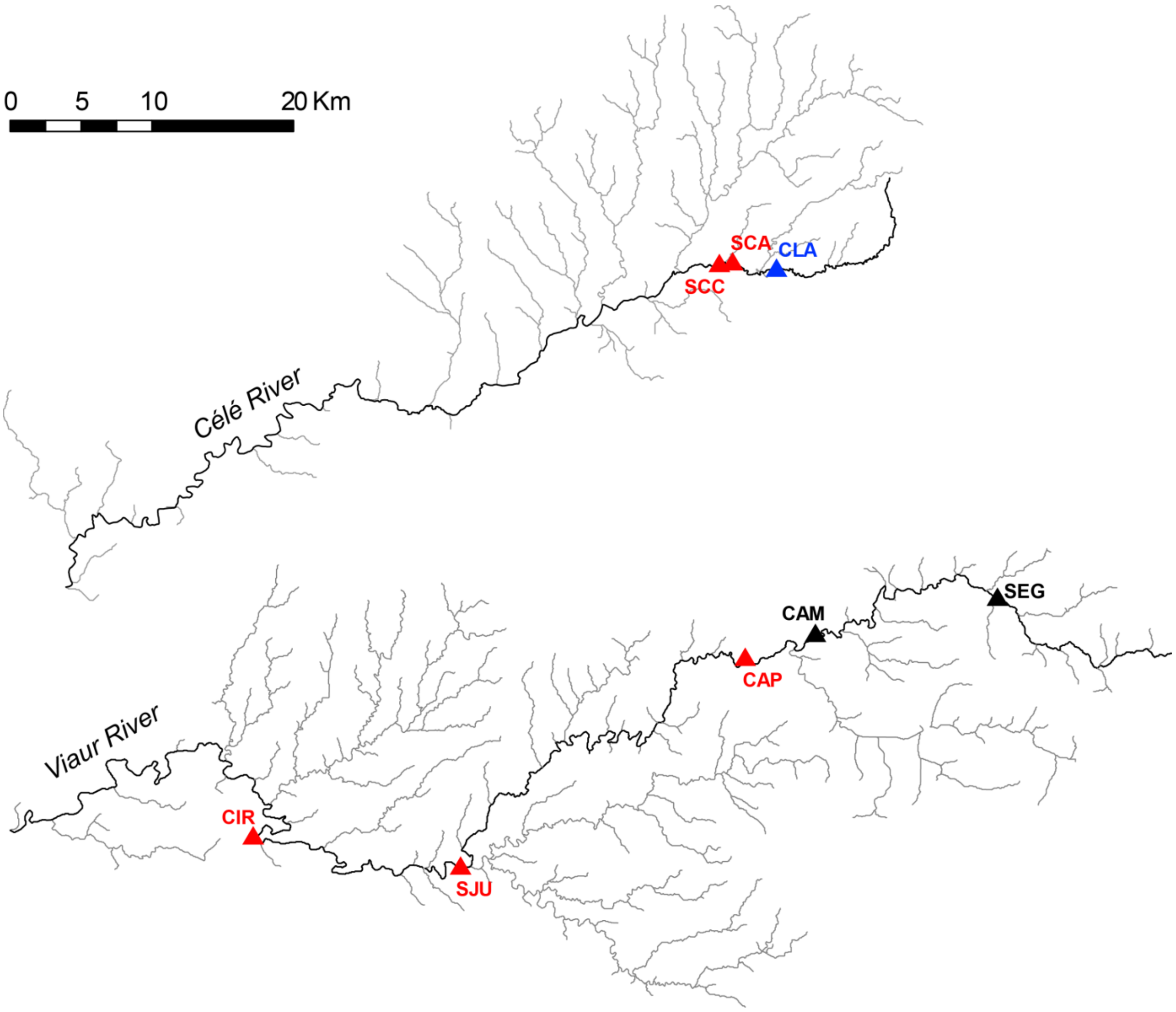

**Appendix S9**

**Figure S9a:** For each mutation rate ($\mu$ = $5.10^{-5}$ or $5.10^{-4}$) and each metric of genetic differentiation (G''st and ϕ'st), density heatmaps displaying the relationship between observed and predicted measures of genetic differentiation. Datasets characterized by low levels of expected heterozygosity (hexagons with light outline) were associated with higher discrepancy between observed and expected measures of genetic differentiation, that is, higher uncertainty. They yet only concerned 3.5% of data. Discarding these data had no consequence on predicted values, as indicated in Figure S9b.

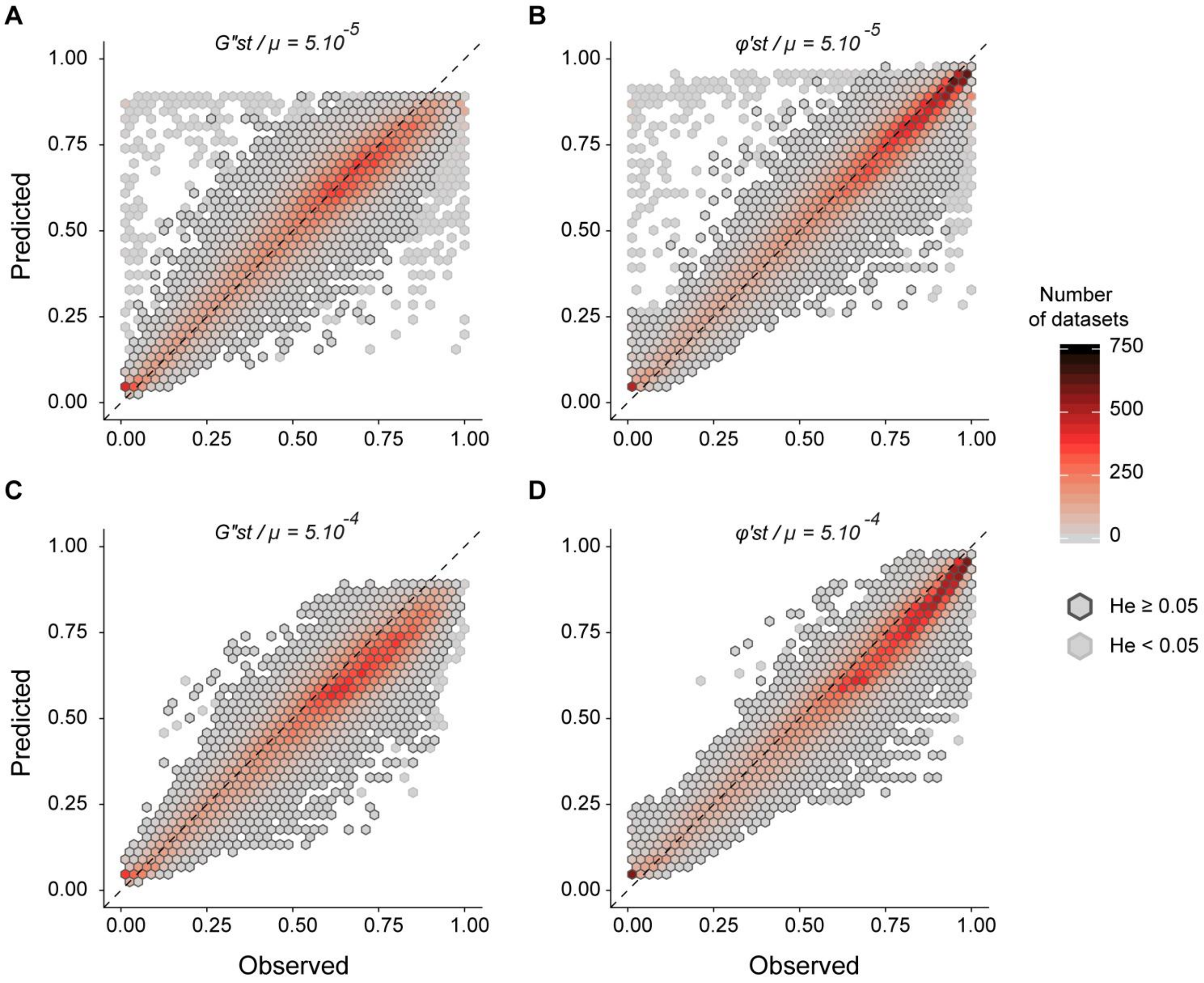

**Figure S9b:** For each mutation rate ($\mu$ = 5.10$^{-5}$ or 5.10$^{-4}$) and each metric of genetic differentiation (G''st and ϕ'st), density heatmaps displaying the relationship between measures of genetic differentiation as predicted from all datasets and measures of genetic differentiation as predicted from a subset of data with *He* ≥ 0.05. Predicted values were highly similar, with Pearson's correlation coefficients higher than 0.999 in all situations.

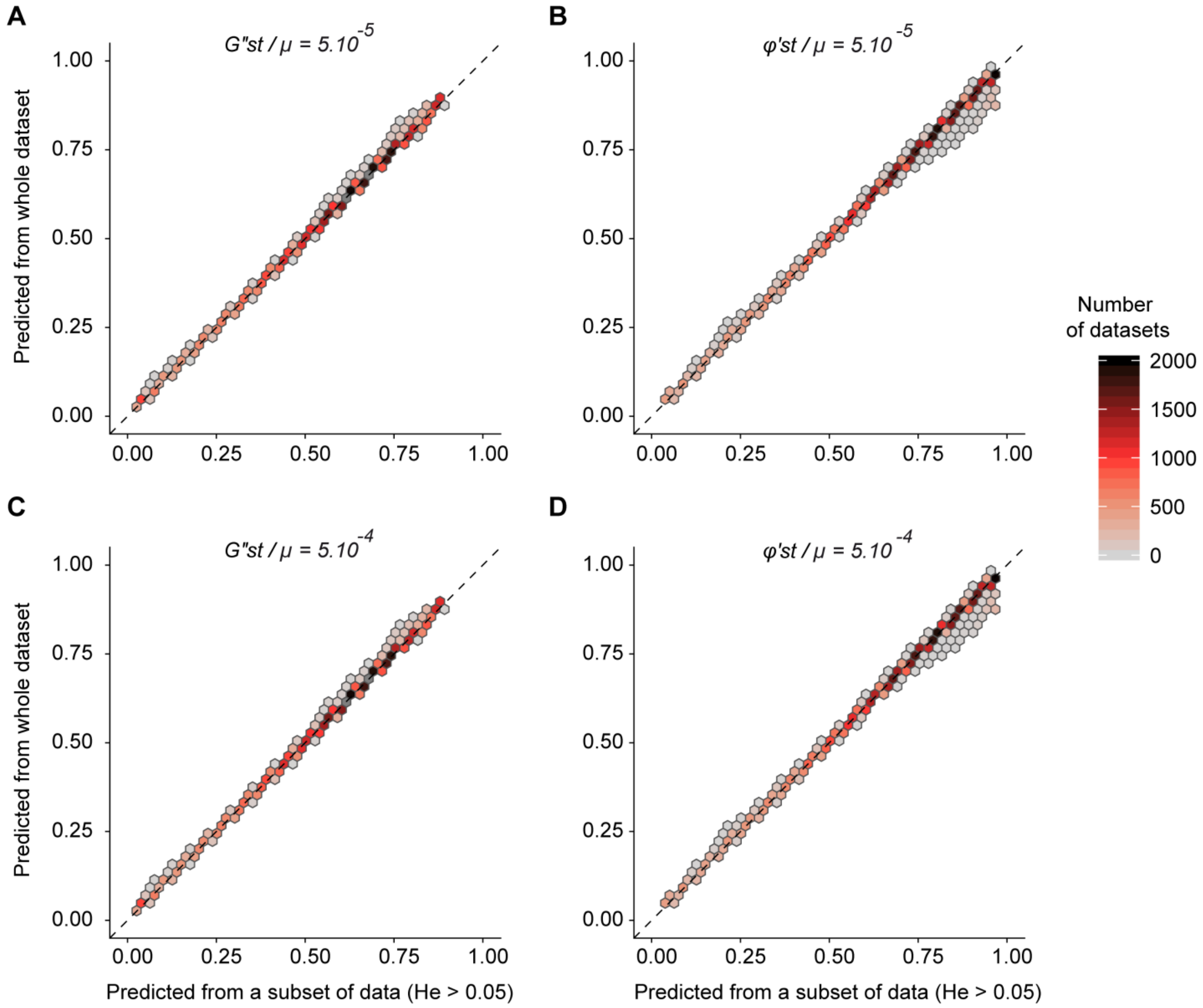

### Appendix S10:

**Figure S10:** For each mutation rate ($\mu$ = 5.10$^{-5}$ or 5.10$^{-4}$) and each metric of genetic differentiation (G''st and ϕ'st), variable importance in RandomForest predictions, expressed as the increase in Mean Square Error following random permutation of each predictor T (number of generations since barrier creation) and He (expected heterozygosity). The G''st metric was more sensitive to T than to He, whereas PhiST was equally sensitive to both predictors, highlighting the relevance of combining these two complementary measures of genetic differentiation in the computation of the $F_{INDEX}$.

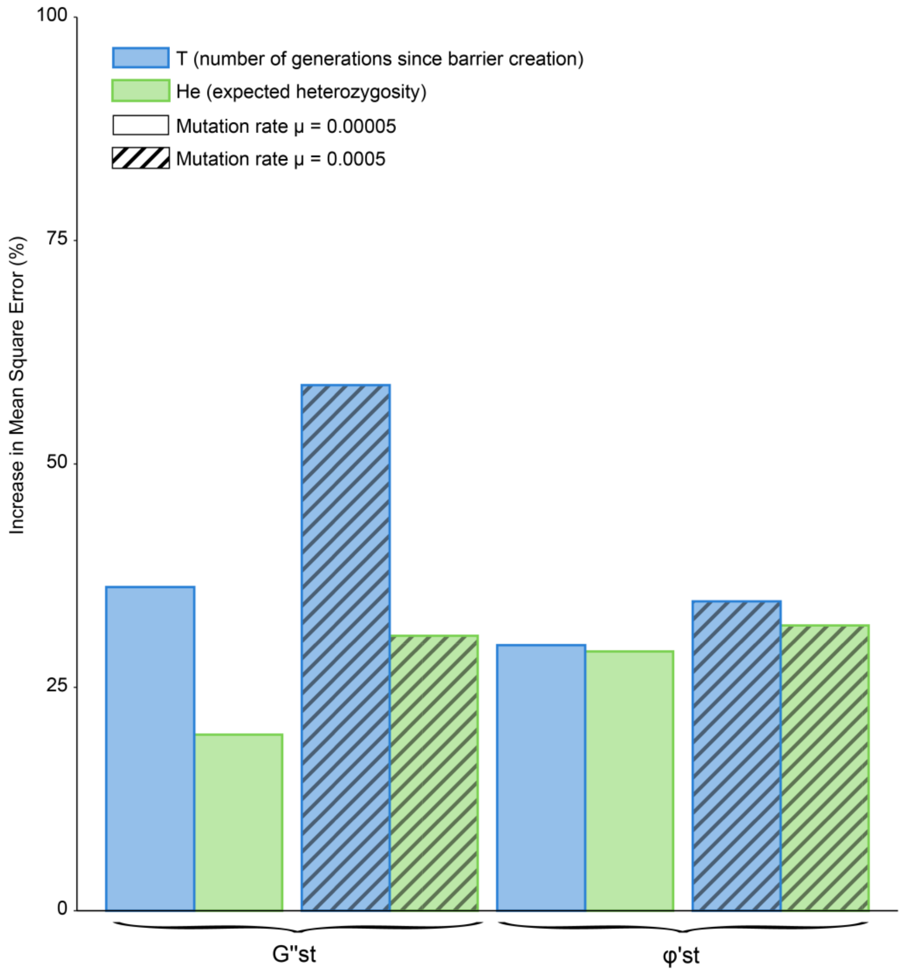

### Appendix S11

In the very first generations after barrier creation, and especially in the case of large populations, the temporal inertia in the evolution of allelic frequencies implies that the difference between $GD_{max}$ and $GD_{min}$ is very low (that is, $GD_{max}$ is also very close from zero). The rescaling performed by the denominator in Equation 1 (see main text and Appendix S2) induces, by construction, an increase in the noise-to-signal ratio in the estimate of $F_{INDEX}$

### Appendix S12

To assess the possible influence of uncertainty in *He* as a proxy for *Ne*, we constructed two subsets from the simulated dataset considered for the theoretical validation of the $F_{INDEX}$. For each combination of K (carrying capacity), *m* (crossing rate) and T (number of generations since barrier creation), we retained simulations from the first ("Low He") and the fourth ("High He") quartiles in terms of *He* values. $F_{INDEX}$ values were computed separately for each subset and plotted against crossing rate, in the same way as the original validation dataset (Figure 3 in main text). Figure S12 illustrates the results for T=100 and for K=50 or 100. Overall, $F_{INDEX}$ values showed high overlap of 95% CI and pointed to the same conclusions whatever the level of *He*: significant barriers for a crossing rate < 0.01 ($F_{INDEX}$ > 90%), intermediate barriers for crossing rates < 0.1, and no barrier effect beyond ($F_{INDEX} \leq$ 20%). The 95%CI about $F_{INDEX}$ values thus correctly capture the uncertainty associated with the use of *He* as a proxy for *Ne*.

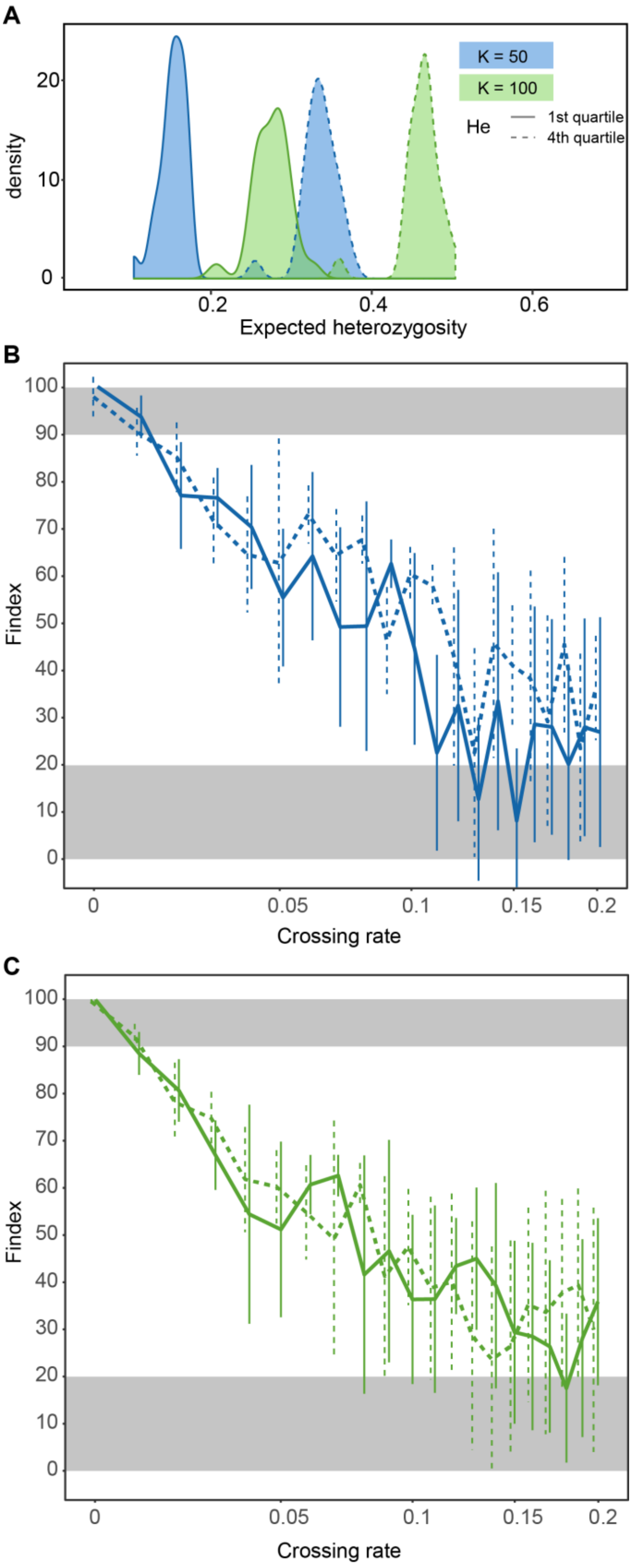


**Figure S12:** (A) For each subset of simulated data (solid lines: 1st quartile / low *He*; dashed lines: 4th quartile / high *He*); density plot of *He* values for K = 50 (blue) and K = 100 (green). (B) $F_{INDEX}$ response to the increase in crossing rate (on a logarithmic scale) for K = 50 and T = 100 generations after barrier creation, according to *He* levels (solid: low *He*, dashed: high *He*). (C) Same as B, but for K = 100.

### Appendix S13

To assess the influence of uncertainty in the number $T$ of generations since barrier creation (stemming from uncertainty in the age of the obstacle and/or in the generation time of the focal species), we used the simulated dataset considered for the theoretical validation of the $F_{INDEX}$. For each of the 21600 simulated genetic datasets, we computed new $F_{INDEX}$ values using a new number $T'$ of generations randomly picked from a uniform distribution ranging from $T$ to $T$+(2×$T$) to mimic an overestimation of $T$ (up to 200%) and from a uniform distribution ranging from $T$-(0.95×$T$) to $T$ to mimic an underestimation of $T$ (up to 95%; we did not consider anunderestimation of 100% as it is usually known whether an obstacle was just built or not in the real world). For an obstacle of age $T$ = 300, $T'$ thus ranged from 15 to 600; for $T$ = 5, $T'$ ranged from 0.25 to 10. We then compared these new $F_{INDEX}$ values with original ones (that is, computed using the number $T$ of generations actually used in simulations) in the form of a $\Delta F_{INDEX}$ and plotted the results against uncertainty (expressed in percentage).

Figure S13 indicates that the $F_{INDEX}$ is almost insensitive to a ~30% underestimation and to a ~50% overestimation of the actual number of generations since barrier creation ($|\Delta F_{INDEX} \leq 5|$): for a barrier of age $T$ = 300, it means that the $F_{INDEX}$ would not show any deviation higher than 5% if the barrier was given an erroneous age $T'$ of only 210 generations or on the contrary of up to 450 generations. Figure S13 further indicates that the $F_{INDEX}$ can be considered as highly robust to a ~60% underestimation and to a ~150% overestimation ($|\Delta F_{INDEX} \leq 10|$): for a barrier of age $T$ = 300, it means that the $F_{INDEX}$ would not show any deviation higher than 10% if the barrier was given an erroneous age $T'$ of only 120 or on the contrary of up to 750 generations. As a rule of thumb, we can thus consider that the $F_{INDEX}$ is highly robust to a ~50% uncertainty in the actual age of the barrier, and that it is more conservative to slightly overestimate the age of the barrier when it cannot be known for sure.

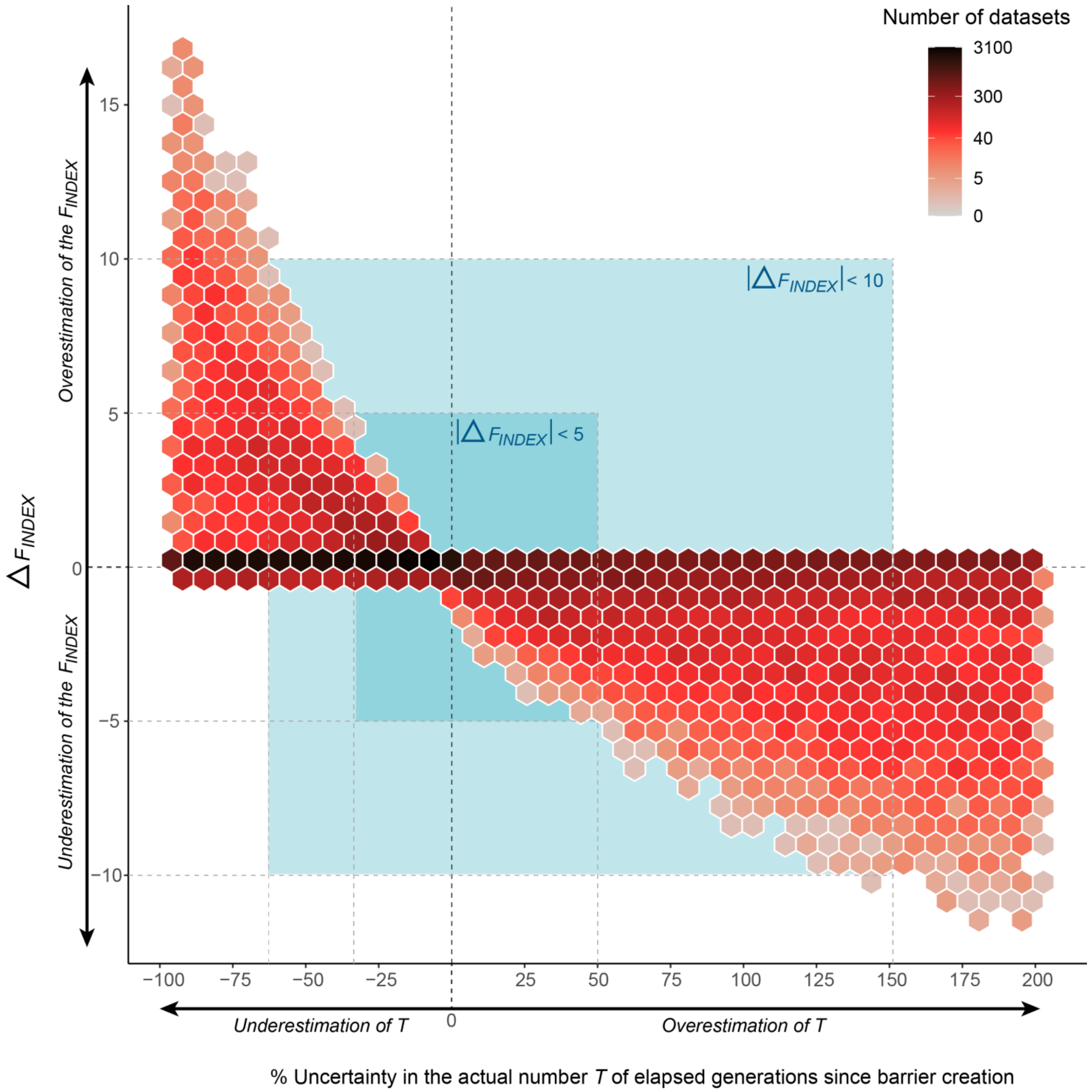


**Figure S13:** Density heatmap of $\Delta F_{INDEX}$ (the difference between $F_{INDEX}$ values computed from an erroneous age of the barrier and $F_{INDEX}$ values computed from the actual age of the barrier) against levels of uncertainty in the age of the barrier (expressed in percentage). Light blue surfaces indicate the parameter space corresponding to $|\Delta F_{INDEX} \leq 5|$ (inner surface) and $(|\Delta F_{INDEX} \leq 10|$ (outer surface). Note the logarithmic scale for the colors of hexagonal cells ('Number of datasets').

### REFERENCES


Adamack, A. T., & Gruber, B. (2014). PopGenReport : Simplifying basic population genetic analyses in R. *Methods in Ecology and Evolution*, *5*(4), 384‑387.

Borenstein, M. (Éd.). (2009). *Introduction to meta-analysis*. John Wiley & Sons.

Bowcock, A. M., Ruizlinares, A., Tomfohrde, J., Minch, E., Kidd, J. R., & Cavallisforza, L. L. (1994). High-resolution of human evolutionary trees with polymorphic microsatellites. *Nature*, *368*(6470), 455‑457.

Cavalli-Sforza, L. L., & Edwards, A. W. (1967). Phylogenetic analysis. Models and estimation procedures. *American journal of human genetics*, *19*, 233.

Goudet, J. (1995). FSTAT (Version 1.2) : A Computer Program to Calculate F-Statistics. *Journal of Heredity*, *86*(6), 485‑486. https://doi.org/10.1093/oxfordjournals.jhered.a111627

Goudet, J. (2005). Hierfstat, a package for r to compute and test hierarchical F-statistics. *Molecular Ecology Notes*, *5*(1), 184‑186. https://doi.org/10.1111/j.1471-8286.2004.00828.x

Gouskov, A., Reyes, M., Wirthner-Bitterlin, L., & Vorburger, C. (2016). Fish population genetic structure shaped by hydroelectric power plants in the upper Rhine catchment. *Evolutionary Applications*, *9*(2), 394‑408. https://doi.org/10.1111/eva.12339

Hague, M. T. J., & Routman, E. J. (2016). Does population size affect genetic diversity? A test with sympatric lizard species. *Heredity*, *116*(1), 92–98. https://doi.org/10.1038/hdy.2015.76

Hedrick, P. W. (2005). A Standardized Genetic Differentiation Measure. *Evolution*, *59*(8), 1633‑1638. https://doi.org/10.1111/j.0014-3820.2005.tb01814.x

Jombart, T. (2008). adegenet : A R package for the multivariate analysis of genetic markers. *Bioinformatics*, *24*(11), 1403‑1405. https://doi.org/10.1093/bioinformatics/btn129

Jost, L. (2008). $G_{\mathrm{ST}}$ and its relatives do not measure differentiation. *Molecular Ecology*, *17*(18), 4015‑4026. https://doi.org/10.1111/j.1365-294X.2008.03887.x

Meirmans, P. G. (2006). Using the Amova Framework to Estimate a Standardized Genetic Differentiation Measure. *Evolution*, *60*(11), 2399‑2402. https://doi.org/10.1111/j.0014-3820.2006.tb01874.x

Meirmans, P. G., & Hedrick, P. W. (2011). Assessing population structure : FST and related measures. *Molecular Ecology Resources*, *11*(1), 5‑18. https://doi.org/10.1111/j.1755-0998.2010.02927.x

Nei, M. (1973a). Analysis of gene diversity in subdivided populations. *Proceedings of the National Academy of Sciences*, *70*(12), 3321–3323.

Nei, M. (1973b). The theory and estimation of genetic distance. In *Genetic Structure of Populations* (University Press of Hawaii, p. 45‑54). Morton N. E.

Nei, M., & Chesser, R. K. (1983). Estimation of fixation indices and gene diversities. *Annals of Human Genetics*, *47*(3), 253‑259. https://doi.org/10.1111/j.1469-1809.1983.tb00993.x

Neuenschwander, S., Michaud, F., & Goudet, J. (2019). QuantiNemo 2 : A Swiss knife to simulate complex demographic and genetic scenarios, forward and backward in time. *Bioinformatics*, *35*(5), 886‑888. https://doi.org/10.1093/bioinformatics/bty737

Prevosti, A., Ocaña, J., & Alonso, G. (1975). Distances between populations of *Drosophila subobscura* , based on chromosome arrangement frequencies. *Theoretical and Applied Genetics*, *45*(6), 231‑241. https://doi.org/10.1007/BF00831894

Prunier, J. G., Dubut, V., Chikhi, L., & Blanchet, S. (2017). Contribution of spatial heterogeneity in effective population sizes to the variance in pairwise measures of genetic differentiation. *Methods in Ecology and Evolution*, *8*(12), 1866–1877. https://doi.org/10.1111/2041-210X.12820

Prunier, J. G., Dubut, V., Loot, G., Tudesque, L., & Blanchet, S. (2018). The relative contribution of river network structure and anthropogenic stressors to spatial patterns of genetic diversity in two freshwater fishes : A multiple-stressors approach. *Freshwater Biology*, *63*(1), 6‑21. https://doi.org/10.1111/fwb.13034

R Development Core Team. (2014). *R: A Language and Environment for Statistical Computing, R Foundation for Statistical Computing*. http://www.R-project.org

Rogers, J. S. (1972). Measures of genetic similarity and genetic distance. In *Studies in Genetics VII* (University of Texas Publication 7213, p. 145‑153).

Rousset, F. (2008). GENEPOP ′007 : A complete re-implementation of the GENEPOP software for Windows and Linux. *Molecular Ecology Resources*, *8*(1), 103‑106.

Sanghvi, L. D. (1953). Comparison of genetical and morphological methods for a study of biological differences. *American Journal of Physical Anthropology*, *11*(3), 385‑404. https://doi.org/10.1002/ajpa.1330110313

Weir, B. S., & Cockerham, C. C. (1984). Estimating F-Statistics for the analysis of population structure. *Evolution*, *38*(6), 1358‑1370.

Winter, D. J. (2012). MMOD : An R library for the calculation of population differentiation statistics. *Molecular Ecology Resources*, *12*(6), 1158‑1160. https://doi.org/10.1111/j.1755-0998.2012.03174.x
